# Weak effects of local prey density and spatial overlap on predation intensity in a temperate marine ecosystem

**DOI:** 10.1101/2025.03.27.645454

**Authors:** Max Lindmark, Christopher A. Griffiths, Valerio Bartolino, Viktor Thunell, Federico Maioli, Sean C. Anderson, Andrea Belgrano, Michele Casini, Katarzyna Nadolna-Ałtyn, Joanna Pawlak, Marzenna Pachur, Marcin Rakowski, Karolina Wikström, Murray S. A. Thompson, Mayya Gogina, Didzis Ustups, Nis S. Jacobsen

## Abstract

Quantifying the impact of lower trophic level species abundance on higher trophic level predators (and vice versa) is critical for understanding marine ecosystem dynamics and for implementing ecosystem-based management. Trophic ecosystem models generally predict a tight coupling between prey and fish predators, such that higher abundance of lower trophic species increases abundance of higher trophic level predators. This assumes that predator feeding rates are limited by prey availability to some degree. Despite being a key component of predator-prey interactions and multi-species fisheries management, relatively few studies have assessed the impacts of prey availability on predation patterns of mobile, generalist fish predators using spatiotemporal models and local-scale stomach content, predator, and prey data. In this study, we explore the association between local density of key prey and predator stomach contents, and predator-prey spatiotemporal overlap and predation indices, using the Baltic Sea as a case study. We use three decades of spatially resolved biomass and stomach content data on Atlantic cod (*Gadus morhua*), and biomass data on three of its key prey: herring (*Clupea harengus*), the isopod *Saduria entomon*, and sprat (*Sprattus sprattus*). Using geostatistical generalized linear mixed effects models fitted to relative biomass density and prey-weight-per-predator-weight, we estimate spatiotemporal trends and annual indices of biomass weighted and area-expanded per-capita and population-level predation, predator-prey overlap, and the correlation between these indices. Range shifts have resulted in reduced predator-prey overlap over time, which is now the lowest in three decades. For *Saduria*, we find an association between prey availability and stomach contents, but not for herring or sprat. Similarly, only in *Saduria* do we find a positive correlation between population-level predation indices and the spatiotemporal overlap. Although behavioral interactions with pelagic prey are challenging to infer from stomach content and acoustic data due to high mobility leading to fine-scale spatiotemporal mismatch, the weak connection with local-scale availability, and low correlation between population-level predation and spatial overlap, could imply weaker coupling between pelagic prey and cod than previously thought. These findings provide key information on the strength of species interactions, which is crucial for the continued development of multispecies models and ecosystem-based fisheries management.

## Introduction

Temperate marine ecosystems are shaped by complex combinations of bottom-up and top-down processes (Lynam *et al*. 2017). In coastal or upwelling systems, these two processes can be linked at intermediate trophic levels by small pelagic planktivorous fish, which can control lower trophic levels with top down effects and upper trophic levels via bottom up effects — also referred to as wasp-waist control (Cury *et al*. 2000, Bakun 2006). Ecological theory generally suggests a tight coupling between prey and predators (Poggiale 1998) such that higher prey densities benefit predator populations. This positive link between prey abundance and predator performance has been supported by a range of trophic ecosystem models (Smith *et al*. 2011, Pikitch *et al*. 2014, Chagaris *et al*. 2020), although this may vary depending on structural model assumptions (Walters *et al*. 2016). Such findings have led in some circumstances to management strategies that aim to reducing fishing pressure on lower trophic level fish to avoid negative impacts on higher trophic levels (Cury *et al*. 2011, Pikitch *et al*. 2012, Chagaris *et al*. 2020).

Empirical support for this form of bottom-up or “donor effect” (i.e., positive effects of prey on the predator but no clear negative effects of predators on prey, *sensu* Jennings and Kaiser 1998) from intermediate to upper trophic levels is, however, fairly limited (Jensen *et al*. 2012). This could be because marine predatory fish are typically mobile generalists, with diets changing through their ontogeny, which implies many but weak trophic links (Jennings and Kaiser 1998, Strong 1992). Not surprisingly, one of the clearer examples of a donor effect is the decline in productivity of Atlantic cod (*Gadus morhua*) in the Barents Sea. This decline occurred because cod had limited ability to switch prey due to low diversity in the intermediate trophic levels, leading to reduced productivity after the main prey species, capelin and herring, collapsed (Mehl and Sunnanå 1991, Hamre 1994, Jensen *et al*. 2012).

Another reason why these effects are difficult to detect relates to spatial dynamics and the scale at which trophic interactions occur (Hunsicker *et al*. 2011). Often trophic interactions are assessed by relating time series of population abundance of predators and prey (Jennings and Kaiser 1998, Overholtz and Link 2007, Hilborn *et al*. 2017), which neglects the important spatial dimension. While there are several examples of the importance of local scale prey availability and donor effects on upper trophic levels, most are for central place foragers (e.g., for seabirds, Crawford 2007, Cury *et al*. 2011, Robinson *et al*. 2015, Hentati-Sundberg *et al*. 2021), and few involve commercially exploited mobile fish predators (Hilborn *et al*. 2017, Pikitch *et al*. 2018) (but see Fall *et al*. 2021). Recent studies have illustrated the potential of spatiotemporal modelling of predator and prey density and stomach content data to quantify fine-scale spatial and temporal variability in diet, and to derive model-based indices of consumption (Grüss *et al*. 2020, Goodman *et al*. 2022, Gartland and Latour 2024). For example, Gartland and Latour (2024) related model-based consumption-indices derived from spatiotemporal models to stock-level biomass of prey and found positive associations. In a similar study, Goodman *et al*. (2022) correlated annual spatial overlap and predation indices, and found that the support for such correlations varied across species. These studies did not, however, include spatially explicit prey covariates, which may be important, because local prey densities can be high and sufficient for predators even though total prey population abundance is low (Hilborn *et al*. 2017), and because the spatial overlap between predators and their prey can vary in the studied time period.

In this study, we investigated the relationship between local prey availability and the relative weight of prey in predator stomachs, as well as the relationship between spatial overlap and predation at a population level, using the Baltic Sea as a case study. We used three decades of biomass and stomach content data from surveys for the predator Atlantic cod (*Gadus morhua*), and biomass data for some of its main prey, sprat (*Sprattus sprattus*), herring (*Clupea harengus*), and the benthic isopod *Saduria entomon* (henceforth only *Saduria*). The central Baltic Sea ecosystem is a species-poor ecosystem where sprat and herring make up more than 75% of the diet by weight in cod around 35 cm, and there is no alternative pelagic prey for cod present in the system (Kulatska *et al*. 2019, Lindmark *et al*. 2025). Moreover, within this time period, the feeding rates on sprat and *Saduria*, as well as the growth, condition and size-at-maturity of cod have declined substantially (Neuenfeldt *et al*. 2020, Mion *et al*. 2021, Lindmark *et al*. 2023, Svedäng *et al*. 2024). Together with an increased natural mortality (ICES 2022a, Eero *et al*. 2023), this has severely impacted the conservation status of cod, to the degree that even in the absence of a commercial targeted fishery (by EU countries), the stock is expected to remain below its biological limit reference point in the near future (ICES 2021). Hence, it is critically important to understand the drivers behind these changes in the physiological performance of cod in the Baltic Sea. Reduced feeding opportunities on benthic and pelagic prey have been proposed as two of the possible underlying causes (Casini *et al*. 2016, Neuenfeldt *et al*. 2020). Reduced feeding on benthic prey is thought to be due to increased competition for dwindling benthic prey resources mainly linked to the deterioration of the benthic habitats (Neuenfeldt *et al*. 2020), while reduced feeding on pelagic prey has been hypothesized to be due to a reduction of the prey in the main areas of cod occurrence (Casini *et al*. 2016, ICES 2023). This has led to suggestions that pelagic fisheries should be limited in the current main distribution area of cod to improve growth (Eero *et al*. 2012, ICES 2015, Casini *et al*. 2016). However, it is largely unknown how the overall local-scale overlap and encounter rates have changed over time, and how the local prey availability affects feeding opportunities for cod.

Here we aim to answer the following research questions: 1) Is the relative weight of sprat, herring, and *Saduria* in cod stomachs related to the local availability of these prey? 2) Have the relative weights of, and predation on, these prey in the diet changed over space and time? 3) Has the spatial overlap between cod and its prey changed over time? and 4) Do spatial overlap indices correlate with population-level predation indices over time?

## Methods

### Data Stomach data

Stomach content data for Baltic cod have been collected by national institutes (opportunistically or within designated programs) over the past decades. In this study, we use mainly data collated in EU funded projects (Huwer *et al*. 2014, Jacobsen *et al*. 2023) that are available at the newly updated International Council for the Exploration of the Sea (ICES) stomach content database (https://www.ices.dk/data/data-portals/Pages/Stomach-content.aspx). We complement these data with data collated in Lindmark *et al*. (2025), and historical data (Huwer *et al*. 2014). We used 31 years (1993–2023) of data (33,243 individual predators), mainly collected on the bi-annual Baltic International Trawl Survey (BITS) conducted in the first and fourth quarter of the year, but also from other cruises conducted in other times of the year (see Huwer *et al*. (2014) and Jacobsen *et al*. (2023) for a detailed description of the data sources). Therefore, the spatial coverage of stomach sampling has varied over time (see Figure 1 for the spatiotemporal distribution of data).

**Figure 1:**
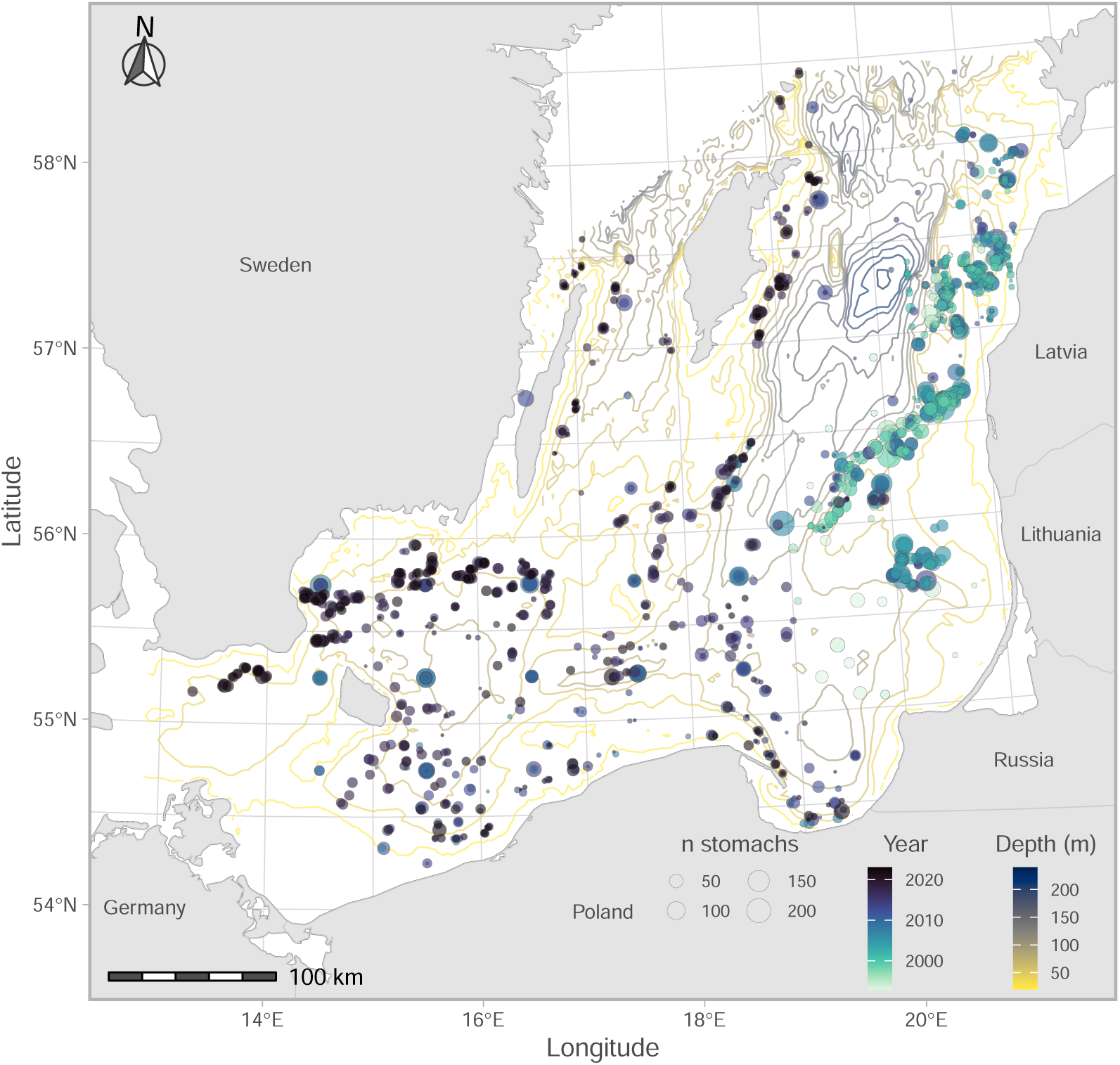
Location of stomach samples in the southern Baltic Sea. Colored contours correspond to depth, the fill color of the points correspond to year, and the size (area) of the points corresponds to sample size by year at that location. Rectangles correspond to ICES rectangles.

Over time, the treatment of regurgitated stomachs (and thereby the classification of empty stomachs) has changed (Neuenfeldt *et al*. 2020). In the early part of the time series, gall bladder status was used to separate non-feeding predators from feeding predators with stomachs regurgitated in the trawl. In recent years, recording gall bladder status is not mandatory, and predators with visible signs of regurgitation are replaced on board. These differences in sampling over time can lead to biased estimates of the proportion of empty stomachs (Neuenfeldt *et al*. 2020). We acknowledge there may be trends due to sampling in the proportion of non-feeding cod and that we cannot assess this bias. Therefore, we opted to include all stomachs in our main analysis, and re-ran the analysis with empty stomachs omitted (across all years, the mean proportion of empty stomachs was 0.19). For each cod predator, we calculated the total weight of sprat, herring and *Saduria* in the stomachs. Any of these prey were found in 43% of predator stomachs. In cases where individual prey length but not prey weight was available, we estimated weight using species specific condition factors and weight-length exponents. The condition factor and exponent for *Saduria* are means for isopods in Robinson *et al*. (2010), and for fish prey (herring and sprat) they were retrieved from FishBase (Froese *et al*. 2010). These individual-level prey weights were used to calculate prey-weight-per-predator-weight — hereafter referred to as “relative prey weight”. The size distribution of cod, and the distribution of relative prey weights can be seen in Appendix S1: Figures S1, S2. Similarly, cod weight was estimated from cod length if missing in the data. Because the length-weight relationship in Baltic cod has varied substantially over the time period, affecting the relative prey weight, we estimated annual values of the condition factor from the trawl survey BITS (see section *Biomass density models*). After data processing, 33,243 cod stomachs were available for analysis.

### Prey data

ICES rectangle-level (Figure 1) biomass estimates for sprat and herring were acquired from the ICES Baltic International Fish Survey Working Group (WGBIFS) database for the Baltic International Acoustic Survey (BIAS) (https://www.ices.dk/community/groups/pages/WGBIFS.aspx). As in Lindmark *et al*. (2023), biomass density of *Saduria* was extracted from a habitat distribution model coupled with modelled hydrographical data from the regional coupled ocean biogeochemical model ERGOM (Gogina *et al*. 2020, Neumann *et al*. 2021). The model was trained to the time period 1981–2019 and predicted for the time period 1993–2019 (but note that this prediction is constant over time and therefore represents the core *Saduria* habitat).

### Predator biomass density

To calculate spatially explicit, weighted predation metrics and predator-prey overlap metrics (Figure 2), we modelled the spatiotemporal distribution of cod using catch per unit effort data (CPUE, numbers/hour) by length class from the fishery-independent Baltic International Trawl Survey (BITS) conducted in the first and fourth quarter between 1993–2023 in the ICES subdivisions 24–28. We used data from the ICES trawl survey database DATRAS (https://www.ices.dk/data/data-portals/Pages/DATRAS.aspx). CPUE data were standardized based on gear dimensions and towing speed (TVL trawl with 75 m sweeps at 3 knots) to units of kg/km^2^, following Orio *et al*. (2017) and Lindmark *et al*. (2023), using length-weight relationships fitted by year to convert from numbers-at-length to weight.

**Figure 2:**
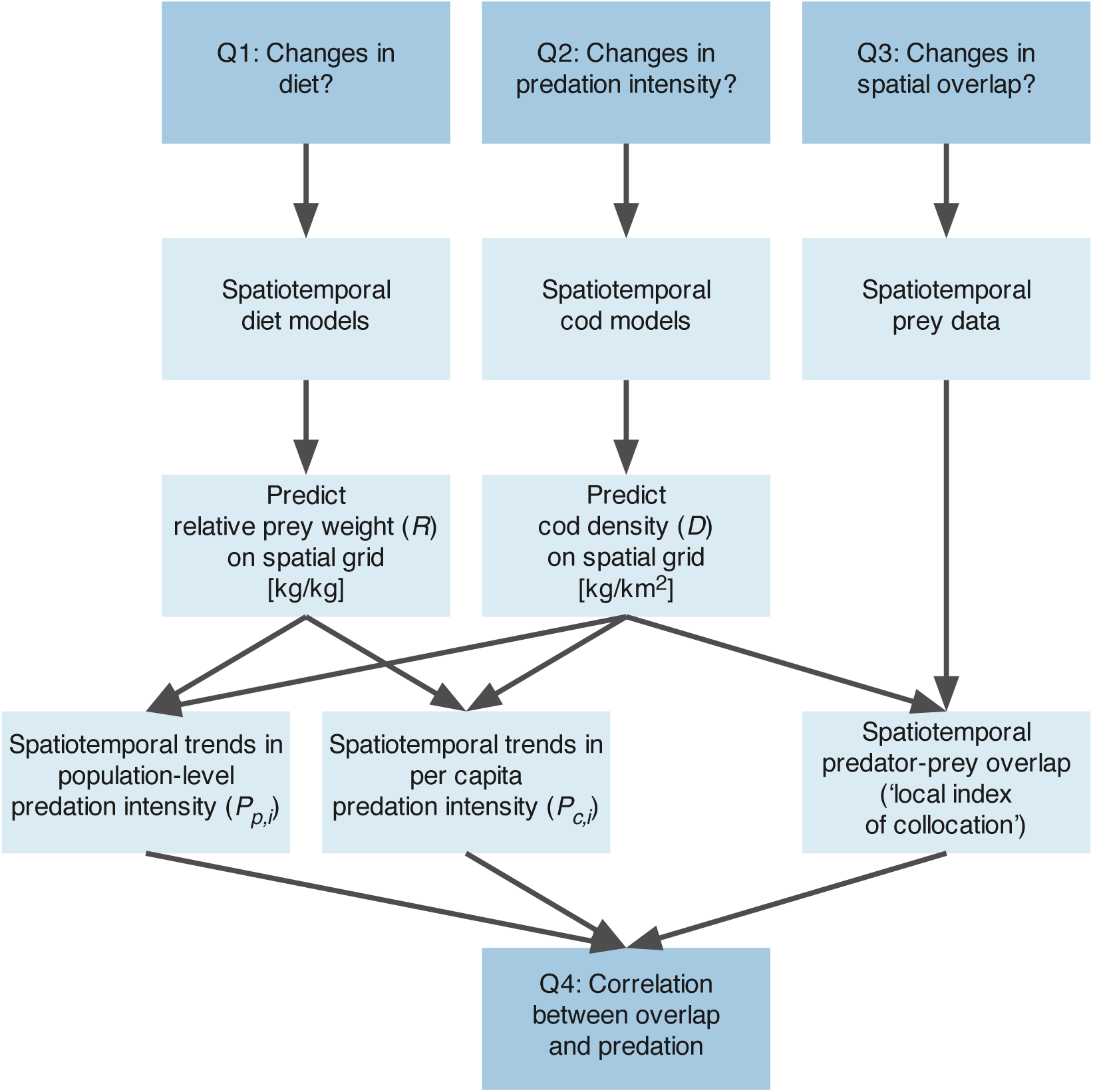
Chart describing the data and modeling process to acquire predation indices (population-level and per-capita) as well as predator-prey overlap. For the definition of *P*_*p*,*i*_ and *P*_*c*,*i*_, see equations 10–11. Research questions are indicated in darker blue boxes.

### Environmental data

We included dissolved sea bottom oxygen concentration (ml/l), sea bottom temperature (℃), and sea bottom salinity (‰) from the Copernicus Marine Service Baltic Monitoring and Forecasting Centre (BAL MFC) as covariates in our biomass density models. Dissolved oxygen values at the sea floor stem from the Baltic Sea biogeochemical model (https://doi.org/10.48670/moi-00009), which is based on ERGOM (Ecological Regional Ocean Model, https://ergom.net/) (Neumann *et al*. 2021), coupled with the NEMO ocean model (Madec *et al*. 2023). Sea floor temperature and salinity stem from the Baltic Sea Physical Reanalysis (https://doi.org/10.48670/moi-00013), based on simulations from the NEMO 3D ocean-ice model version 4.0 (Gurvan *et al*. 2019). These variables were matched to the survey catch data on a monthly resolution. We also included depth (m) in the model, which was was extracted from the EMODnet Bathymetry project (https://emodnet.ec.europa.eu/en/bathymetry) (EMODnet Bathymetry Consortium 2022).

### Spatiotemporal modelling framework

#### Model description

We used spatiotemporal GLMMs (generalized linear mixed effects models) to model stomach contents and biomass density of cod. Per-capita and population-level predation, and predator-prey overlap, were calculated from predictions onto a 3×3 km spatiotemporal grid. Figure 2 illustrates the workflow and how the models and data come together. The full model can be written as follows, but note that the biomass density and diet models do not contain all these terms:

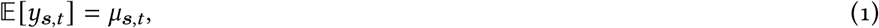

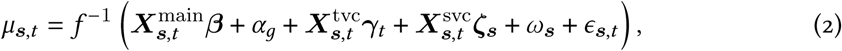

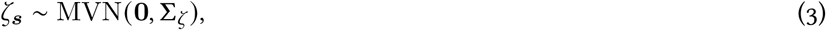

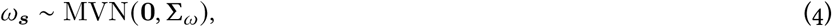

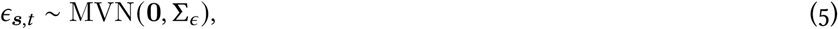

where *y****_s,t_*** is the response variable (relative prey weight or cod biomass density) in location ***s*** at time *t*, *μ* is the mean, *f*^−1^ is the inverse link function, ***X***^main^, ***X***^tvc^, and ***X***^svc^ are design matrices for fixed effects, time-varying coefficients, and spatially varying coefficients, respectively. Their corresponding coefficient vectors are ***β***, ***γ***_*t*_, and ***ζ_s_***. All fixed effect covariates were standardized by subtracting their mean and dividing by their standard deviation. The parameter *α*_*g*_ represents a random intercept for month. The spatially-varying coefficients (*ζ****_s_***), spatial (*ω****_s_***), and spatiotemporal random effects (*ε****_s_***_,*t*_), are assumed drawn from Gaussian Markov random fields (GMRFs) with covariance matrices (i.e., inverse precision matrices) *Σ*_*ζ*_, *Σ*_*ω*_, and *Σ*_*ε*_ constrained by Matérn covariance functions (Rue *et al*. 2009, Lindgren *et al*. 2011). The spatially varying coefficients *ζ****_s_*** are included in the biomass density model to allow the distribution of cod to vary between quarters in addition to the variation given by changes in dynamic covariates, while *ω****_s_*** and *ε****_s_***_,*t*_ reflect spatially correlated latent effects that are constant through time and that vary through time, respectively. The Stochastic Partial Differential Equation (SPDE) approach (Lindgren *et al*. 2011), which links continuous Gaussian random fields with discretely indexed GMRFs, requires piece-wise linear basis functions defined by a triangulated mesh. We defined this mesh using triangles with a cutoff distance (minimum distance between vertices) of 8 km and 10 km for diet and biomass models, respectively, and kept all other arguments in fm_rcdt_2d_inla() (fmesher R package; Lindgren 2023) at their defaults (Appendix S1: Figures S3–S4). Across all models, the cutoff distance was between 2.5–10 times smaller than the estimated Matérn range, which is defined as the distance where spatial correlation effectively disappears (≈ 0.13, Lindgren *et al*. 2011).

To propagate uncertainty in both stomach content predictions and cod density predictions when calculating predation and overlap indices (see sections *Predation indices* and *Spatiotemporal overlap indices*), we simulated 500 draws from the joint parameter precision matrix to make model predictions on the grid. For each draw, we calculated overlap or predation metrics and we present the median, mean, or coefficient of variation of these draws. We predict from the model for a cod length of 33 cm, which is the mean in the diet data.

#### Diet models

The relative prey weight models were fit to each prey species separately. Since these response variables are positive continuous and contain zeroes, we modelled them with either a Tweedie distribution (herring and sprat) or as a “Poisson-link” delta-gamma model (Thorson 2018) (*Saduria*), which has the flexibility of a classic delta or hurdle model (Aitchison 1955), but enables a simpler interpretation of covariates due to log links on both linear predictors (Thorson 2018, Anderson *et al*. 2024). The model family (delta or Tweedie) was chosen based on convergence diagnostics. We included a linear depth effect on all prey, as it has been shown previously to affect cod diets (Pachur and Horbowy 2013). Month was included as a random effect in the *Saduria* model, but not the herring and sprat models because the standard deviation of the month random effect was estimated near zero. We included a linear effect of predator length, reflecting the underlying ontogenetic diet shift cod undergo. To be able to interpolate across a missing year (2011), we modelled the intercept as a time-varying coefficient following an AR(1) process:

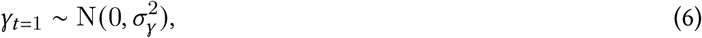

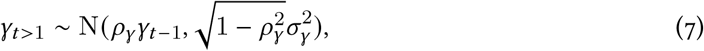

where *ρ* is the correlation between subsequent years and *σ*^2^ is the variance. We included spatially correlated latent effects (*ω****_s_***) that are constant over time and that vary independently each year (*ε****_s_***). Lastly, we included prey biomass or biomass densities as covariates. For the sprat and herring models, this was the estimated biomass per ICES rectangle for ages 1–8+, while for *Saduria*, we extracted rasterized local biomass densities. For each prey, we explored three models: a breakpoint (hockey-stick) function (corresponding to a type I functional response with saturation or a type II functional response), a linear effect, and one without prey availability as covariate. In the break-point term, prey × *b*_0_ is the effect below the threshold, and *b*_0_ × *b*_1_ is the value at the asymptote. We compared the models using the (marginal) Akaike information criterion (AIC; Akaike 1973).

#### Biomass density models

We modelled cod biomass density as a Poisson-link delta-gamma model (Thorson 2018) and included year and quarter as factors, where the latter was in addition modelled as a spatially varying effect, linear effects of salinity, temperature and temperature squared, as well as a breakpoint effect of oxygen, to reflect that dissolved oxygen tends to correlate with biomass density up to a certain point (Essington *et al*. 2022). We also included a time-varying effect of depth and depth squared following a random walk. This to allow the unimodal depth-preferences to change over time (English *et al*. 2022), in line with the shallowing that has been observed for eastern Baltic cod (Orio *et al*. 2019, Lindmark *et al*. 2023).

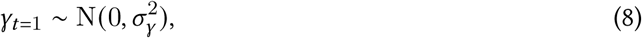

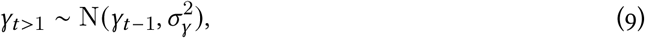

Lastly we included both spatial random effects that are constant in time (*ω****_s_***) and spatial random effects that are independent each year (*ε****_s_***_,*t*_).

#### Model fitting

We fit the spatiotemporal models with the R version 4.3.2 (R Core Team 2024) package sdmTMB (Anderson *et al*. 2024), version 0.6.0.9023. sdmTMB uses automatic differentiation and the Laplace approximation from the R package TMB (Kristensen *et al*. 2016) and sparse matrix structures to set up the SPDE-based Gaussian Markov random fields from the R package fmesher (Lindgren 2023). We estimated parameters via maximum marginal likelihood using the non-linear minimizer nlminb (R Core Team 2024). We confirmed that the optimization was consistent with convergence by checking that the Hessian matrix was positive definite and the maximum absolute log-likelihood gradient with respect to fixed effects was *<*0.001. We evaluated consistency of the models with the data by calculating simulation-based randomized quantile residuals (Dunn and Smyth 1996, Hartig 2022) (Appendix S1: Figures S5–S6). When calculating expected values for the purposes of residual calculation, we took a single draw of the random effects from their multivariate normal distribution (Waagepetersen 2006) rather than using the empirical Bayes random effects estimates. This acknowledges that the random effects are estimated from a distribution (Waagepetersen 2006, Thorson and Kristensen 2024, p. 41).

### Predation indices

We followed the approach presented in Goodman *et al*. (2022) to calculate density-weighted per-capita and population-level predation intensity based on model-predicted relative prey weights and predator densities across the spatiotemporal grid. Population-level predation intensity, *P*_*p*,*i*_, for year *i* was calculated as:

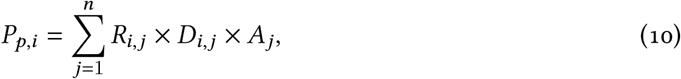

where *R*_*i*,*j*_ is the prey-specific relative prey weights (kg/kg) in grid cell *j*, *D*_*i*,*j*_ is the predicted cod density (kg/km^2^), and *A*_*j*_ is the area of the grid cell (km^2^). This represents a spatially explicit, density-weighted measure of predation intensity (an instantaneous “snapshot” of total weight of a prey species in cod stomachs in units of kg) (Goodman *et al*. 2022). Temporal trends in predation intensity in the domain were acquired by summing grid-level predictions by year. Prior to calculating grid-cell level predation, we omitted grid cells with cod biomass density predictions greater than the 99.99^th^ percentile across all simulations, and relative prey weight predictions *>*1 (less than 0.001% of rows in simulated values of relative sprat weight). These filters did not have a qualitative impact on the calculated metrics, but made sure draws where ecologically realistic. We also omitted areas deeper than 130 m in the prediction grid, since trawl surveys are not conducted at those depths.

Since both cod and pelagic species have undergone shifts in their spatial distribution in this time window (Bartolino *et al*. 2017, Orio *et al*. 2019, Lindmark *et al*. 2023, ICES 2023), and the average feeding rate of cod has declined over time (Neuenfeldt *et al*. 2020), we also wanted to disentangle the effect of distribution shifts from changes in mean feeding rates. This was done by dividing the population-level predation (*P*_*p*,*i*_) by the population-level cod biomass in year *i* (Goodman *et al*. 2022), which yields a weighted average prey biomass per unit predator biomass (per-capita predation), *P*_*c*,*i*_:

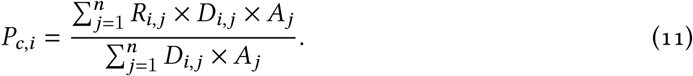

#### Spatiotemporal overlap indices

Predator-prey overlap metrics were calculated from grid-level (3×3 km) predictions of cod biomass density, biomass density of *Saduria* and biomass of sprat and herring at the ICES-rectangle level (Figure 1). There are numerous ways to calculate overlap metrics with slightly different interpretations (for a review, see Carroll *et al*. 2019). In this study, we use the “local index of collocation” overlap metric (Pianka 1973), as in Goodman *et al*. (2022):

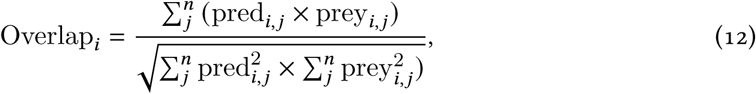

where overlap in year *i* is calculated from the proportions of total biomass of cod (pred_*i*,*j*_) and its prey (prey_*i*,*j*_) in grid cell *j*. We use the same grid as for the stomach content predictions. This metric ranges between 0 and 1 and estimates co-occurrence using correlations between predator and prey densities at the grid-scale, and is suitable for estimating encounter rates (Carroll *et al*. 2019, Goodman *et al*. 2022). When visualizing the overlap in space, we omit the summation across grid cells.

For pelagic prey (sprat and herring), we used only cod predictions in the fourth quarter, since that is when the hydro-acoustic survey (BIAS) takes place. After confirming that the difference between quarters was minimal for overlap with *Saduria* with respect to trends, we presented only results for quarter 4 also for *Saduria*. Given that the spatial predictions of *Saduria* densities are constant over time, changes in overlap with *Saduria* are only driven by changes in the distribution of cod, while in reality, *Saduria* also likely have changed their spatial distribution (Gogina *et al*. 2020).

To quantify how overlap between cod and its prey was related to the predation intensity, we tested the correlation between annual predation intensity (per-capita and population-level) and the annual spatial overlap.

Lastly, to highlight trends over time in per capita and population-level predation, and spatial overlap, we fit generalized additive models (GAMs) to the annual indices, with prey-specific smooth effects of year. We assumed Gamma-distributed errors and a log link for the predation indices, and Beta-distributed errors and a logit link for the overlap indices. The models were fitted with the R package brms (Bürkner 2017). We used default priors, i.e, Student-t(3, 0, 2.5) for the intercept and the standard deviations of spline coefficients, and flat priors for the prey-coefficients. The shape parameter in the Gamma model, and the precision parameter in the Beta model, were given a Gamma(0.01, 0.01) prior. We visualized predictions by summarizing draws from the expectation of the posterior predictive distribution, using the R package tidybayes (Kay 2019).

## Results

The relative weight of a prey species in cod stomachs was positively related to the local density of that prey for *Saduria*, but not for sprat or herring. For sprat, the model without prey biomass was favored by AIC (Table 1). For herring, the breakpoint model and the model without herring were indistinguishable in terms of AIC (Table 1), and the herring covariate was negatively related to herring in the stomachs of cod. Therefore, further analysis were based on the simpler model without the breakpoint herring covariate. The breakpoint model had the lowest AIC among the *Saduria* models, however, the conditional predictions showed high uncertainty on the total prediction (both model components combined) (Appendix S1: Figure S7; Table. S1).

**Table 1:**
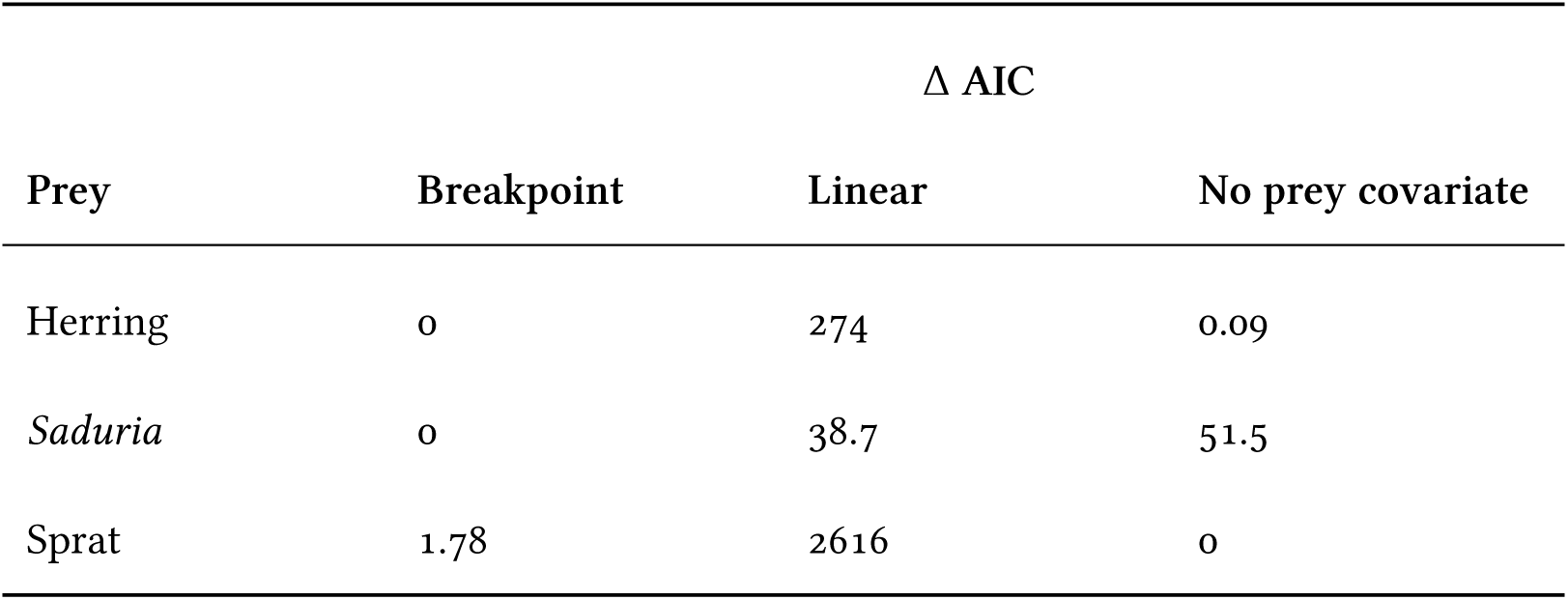
ΔAIC (marginal AIC for the model relative to the model with the lowest AIC) for all models fitted to stomach content data.

We found clear spatial patterns in both stomach contents and predation indices (Figures 3, 4). The relative prey weight of herring in the cod stomachs showed low spatiotemporal variation compared to the other prey apart from very local hotspots from year to year (Figure 3a, d). Both relative prey weight of, and predation on, *Saduria* were highest in the central parts of the southern Baltic Sea (Figures 3b, e, 4b, e), which corresponds to the core area of the *Saduria* distribution in this region (Appendix S1: Figure S8b). Both the relative prey weight of sprat (Figure 3c, f) and the predation on sprat (Figure 4c, f) occurred throughout the Baltic Sea, although the predation was more limited to the southwestern part in recent years due to the shift in distribution of cod (Figure 5).

**Figure 3:**
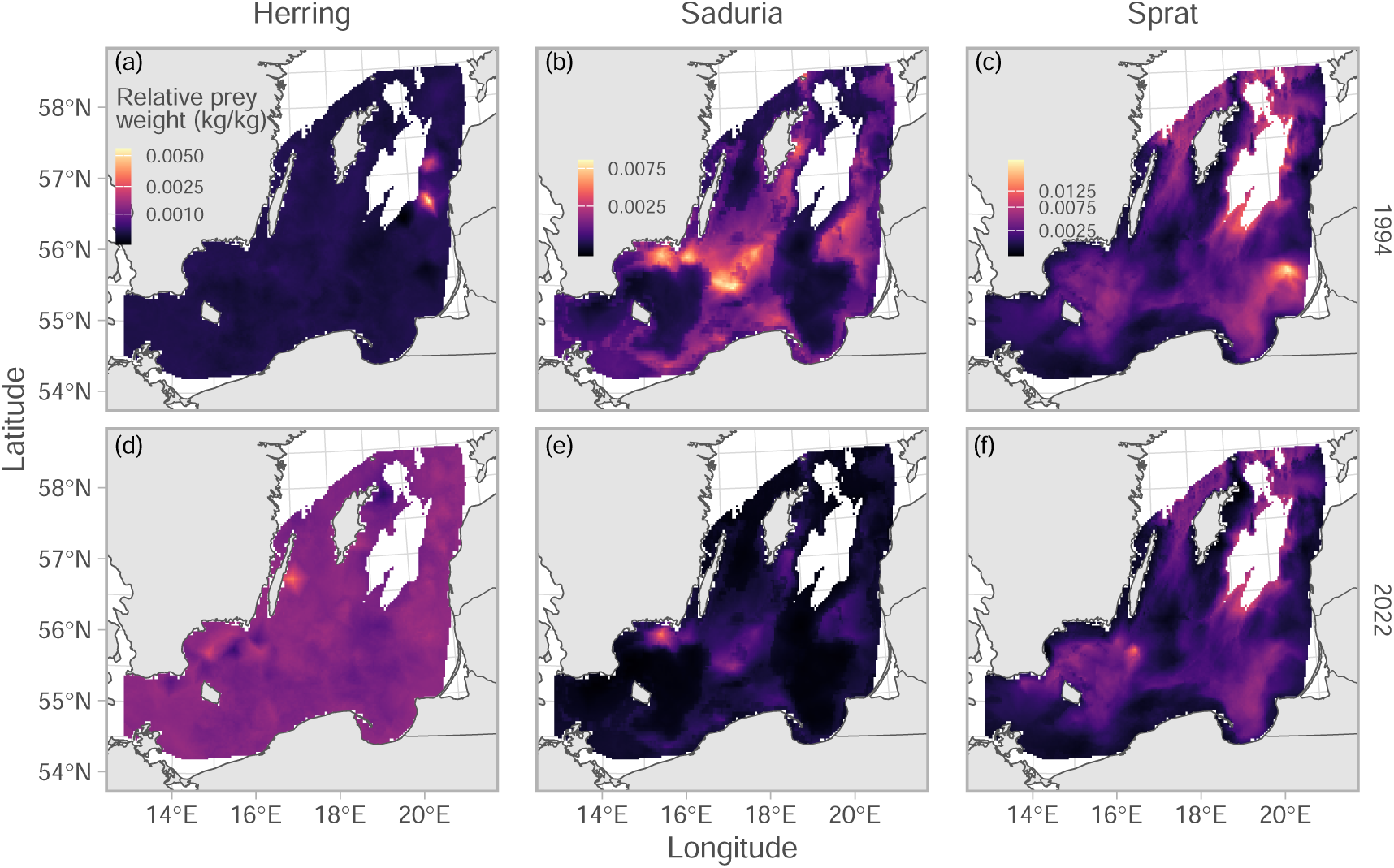
Relative prey weight for a cod of 33 cm. Colors indicate the median value across 500 simulated spatial predictions for herring (a, d), *Saduria* (b, e), and sprat (c, f), for years 1994 (top row, a–c) and 2022 (bottom row, d–f), as examples. The color scale is square-root transformed to better visualize the spatial patterns. Note that scales are shared within species, across years. Only grid cells with depth *<*130 m are included in the plot.

**Figure 4:**
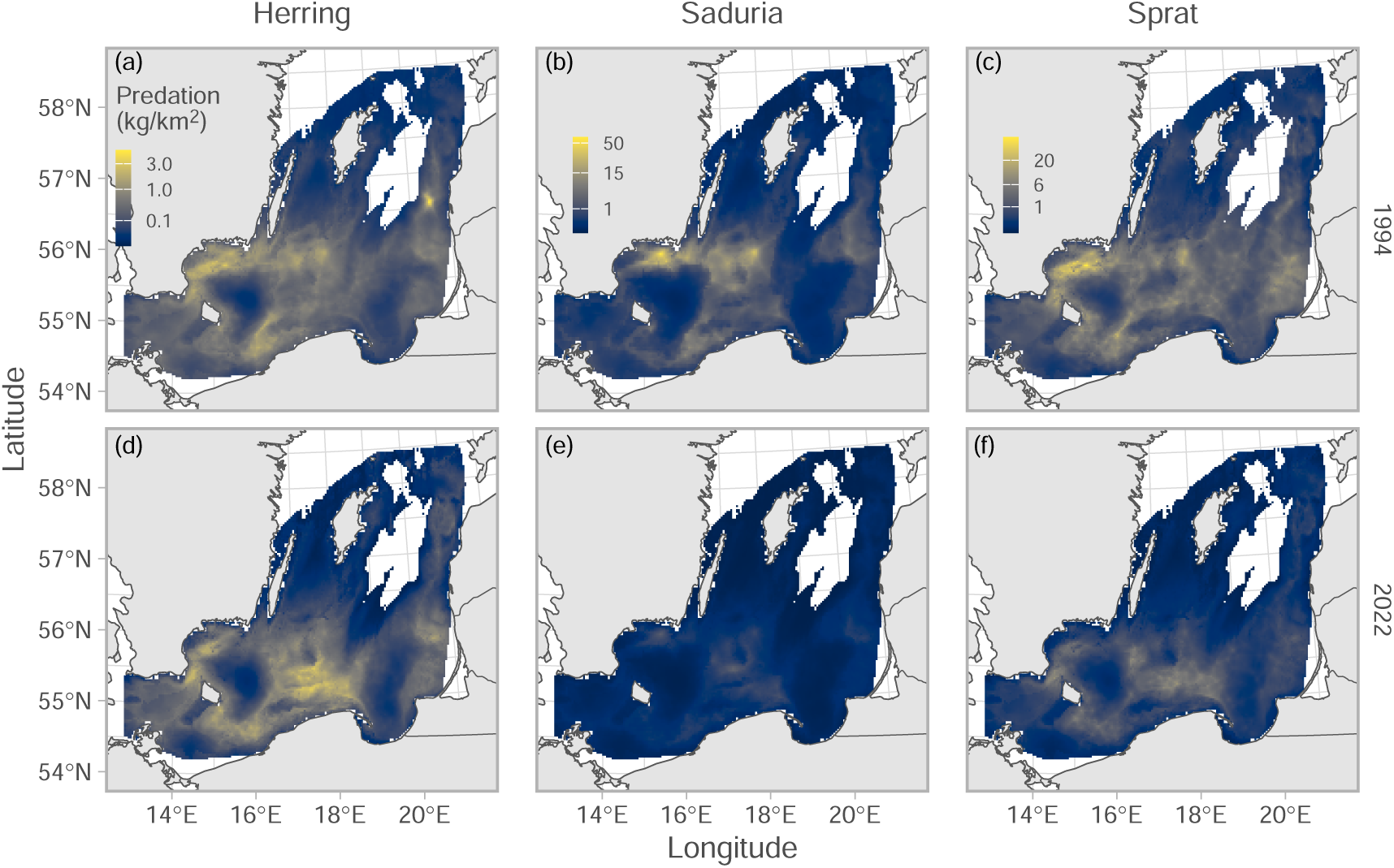
Cod density × relative prey weight (i.e., predation) plotted in space. Colors indicate the mean across 500 simulated spatial predictions of both relative prey weight and cod density for herring (a, d), *Saduria* (b, e), and sprat (c, f), for years 1994 (top row, a–c) and 2022 (bottom row, d–f), as examples. The sum of this metric (expanded by grid cell area) across space is the relative population-level predation intensity depicted in Figure 6. Note that the color scale is 3rd-root transformed, and that scales are shared within species, across years. Only grid cells with depth *<*130 m are included in the plot.

**Figure 5:**
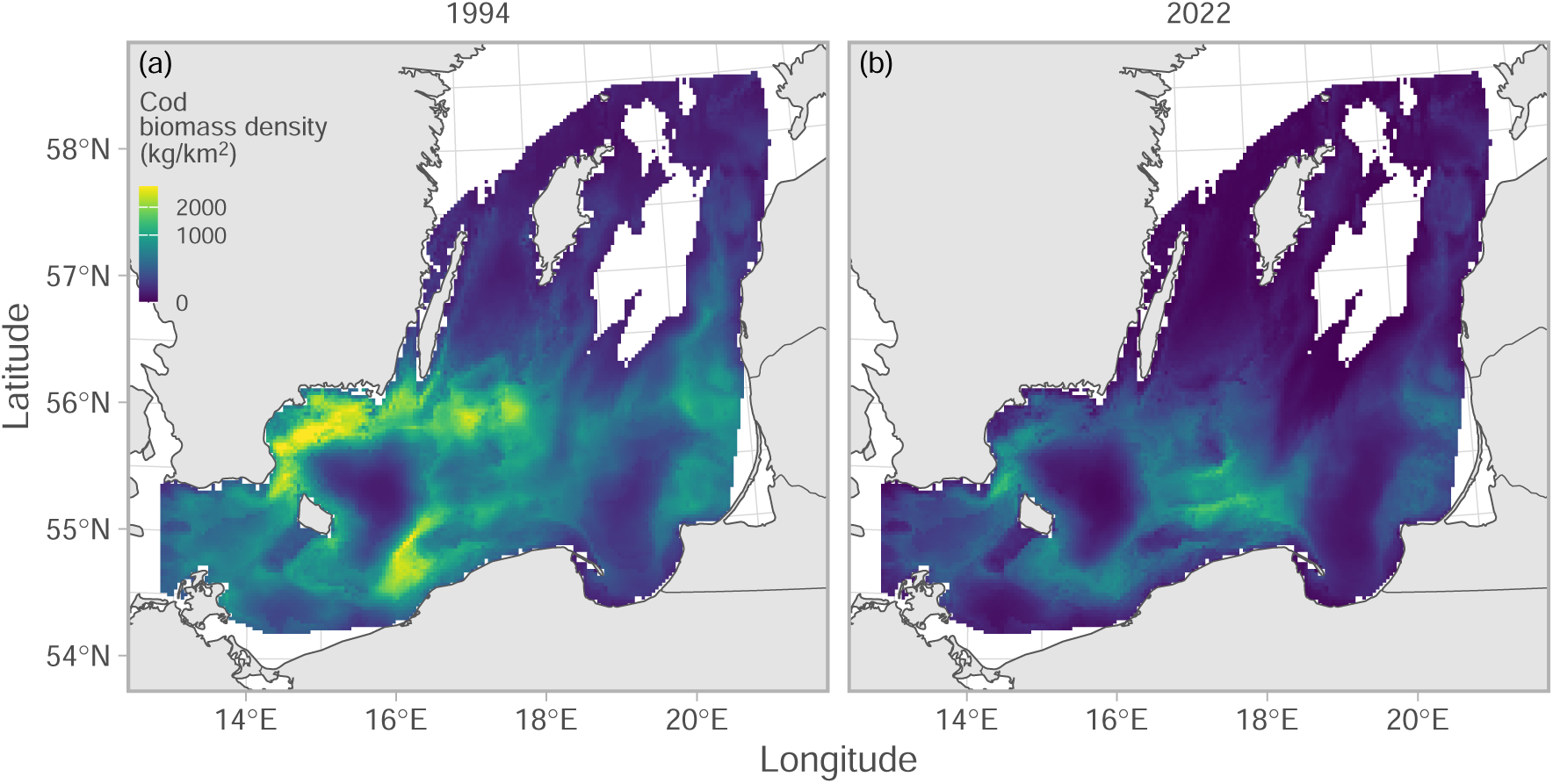
Cod biomass density in space. Colors indicate the mean across 500 simulated spatial predictions for years 1994 (a) and 2022 (b), as examples. Note that the color scale is square-root transformed and truncated at the 99.9^th^ percentile to better visualize the spatial patterns (maximum cod biomass density is 3608 kg/m^2^). Only grid cells with depth *<*130 m are included in the plot.

Over time, the area-expanded per-capita predation on herring increased steadily (with a small decline around the mid 2000s), while the population-level predation reached a peak around 2010 after which it declined to levels similar to the early 2000s (Figure 6a, d). Both per-capita and population-level predation on *Saduria* showed a peak at around 2007, after which it declined steadily to very low levels (Figure 6b, e). Both per-capita and population-level predation on sprat declined in the early 1990s (Figure 6c, f). From then, the per-capita predation on sprat varied some-what cyclically, and showed a weak tendency for an overall increase throughout the time period (Figure 6c). The population-level predation on sprat instead peaked around 2010, and has declined since (Figure 6f). The uncertainty around the per-capita and total predation estimates are substantial when accounting for uncertainty in both cod biomass density and relative prey weights, especially for *Saduria* early in the time series and sprat (Figure 6b, c), and this is largely due to higher uncertainty in the stomach content predictions (Appendix S1: Figure S9). Time series of per-capita and population-level predation were nearly identical for *Saduria* and sprat when empty stomachs were omitted. For herring, trends over time tended to be flattened when empty stomachs where omitted (Appendix S1: Figure S10.

**Figure 6:**
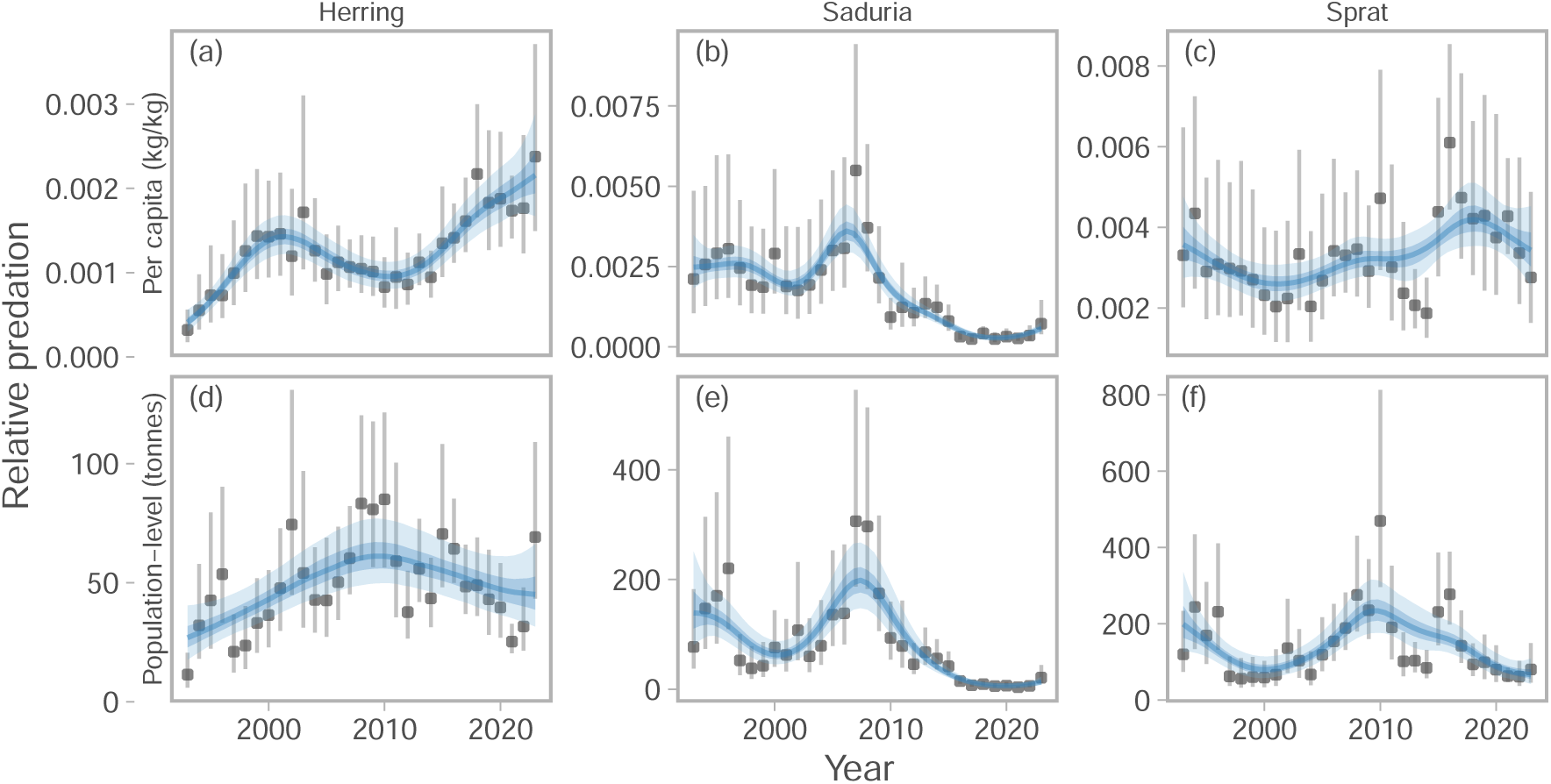
Relative per-capita (top row, a–c) and population-level predation (bottom row, d–f) by cod on herring (a, d), *Saduria* (b, e), and sprat (c, f) over time. Points depict the median predation, and vertical lines depict the range between the 10^th^ and 90^th^ percentile of predation, calculated from 500 simulated spatial predictions of both relative prey weight and cod density. Blue lines depict fits from a generalized additive model with year modelled as a penalized spline, and ribbons correspond to the 50% and 90% credible interval of the prediction.

The overlap between cod and its prey was highest in the central parts of the southern Baltic Sea, and along the southeast coast of Sweden (Figure 7). Over time, the overlap with herring started to decline around 2005 (Figure 8a). The spatial overlap with *Saduria* increased slightly between 1993–2009, but since 2010 it has been lower then average, resulting in a weakly negative trend over time (Figure 8b). The spatial overlap with sprat also declined over time, but in a more cyclic fashion. The current spatial overlap with sprat is the lowest since 2007 (Figure 8c).

**Figure 7:**
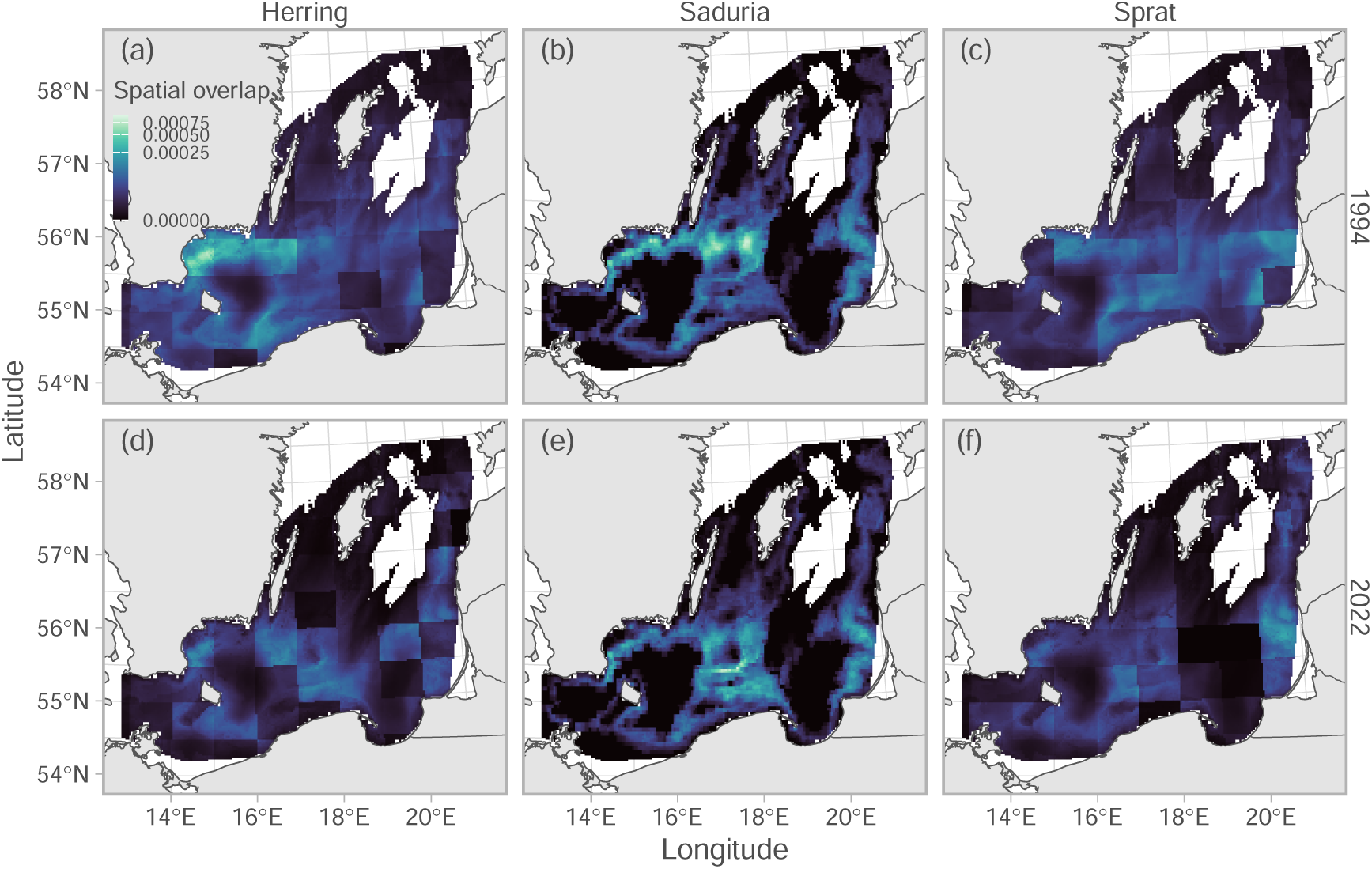
Spatial overlap between cod and its prey herring (a), *Saduria* (b), and sprat (c). Colors indicate the mean across 500 simulated spatial predictions of cod density in years 1994 (top row, a–c) and 2022 (bottom row, d–f), as examples. The color scale is 3rd root transformed to better visualize the spatial patterns. Only grid cells with depth *<*130 m are included in the plot.

**Figure 8:**
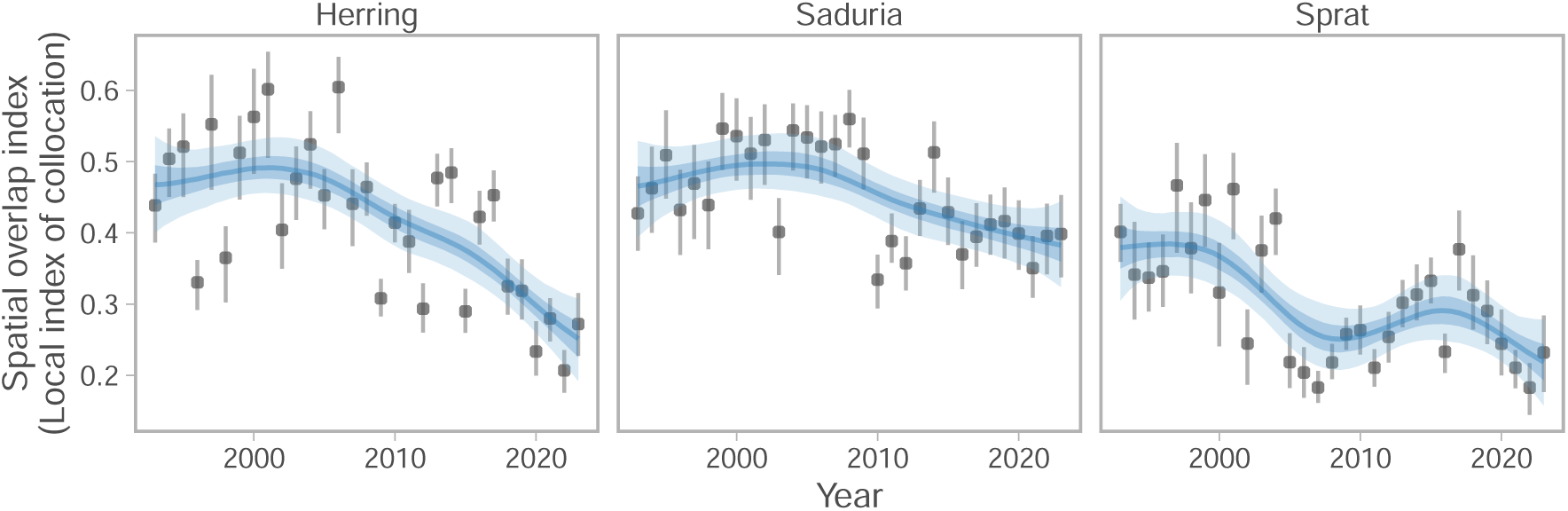
Spatial overlap between cod and its prey herring (a), *Saduria* (b), and sprat (c). Points depict the median overlap, and vertical lines depict the range between the 10^th^ and 90^th^ percentile of overlap, calculated from 500 simulated spatial predictions of cod density. Blue lines depict fits from a generalized additive model with year modelled as a penalized spline, and ribbons correspond to the 50% and 90% credible interval of the prediction.

The correlation between annual estimates of per-capita- and population-level predation with spatial overlap was only clearly positive for *Saduria* (Figure 9b, e). For herring and sprat, the correlation coefficients ranged between −0.39 and −0.12, but the confidence intervals overlapped zero in all cases but the per-capita predation on herring (Figure 9a,c,d,f). The results were nearly identical when fitting diet models to data with empty stomachs omitted for *Saduria* and sprat, whereas for herring the main difference was that the confidence intervals of the correlation between per-capita herring predation and cod-herring overlap crossed 0 (Appendix S1: Figure S11).

**Figure 9:**
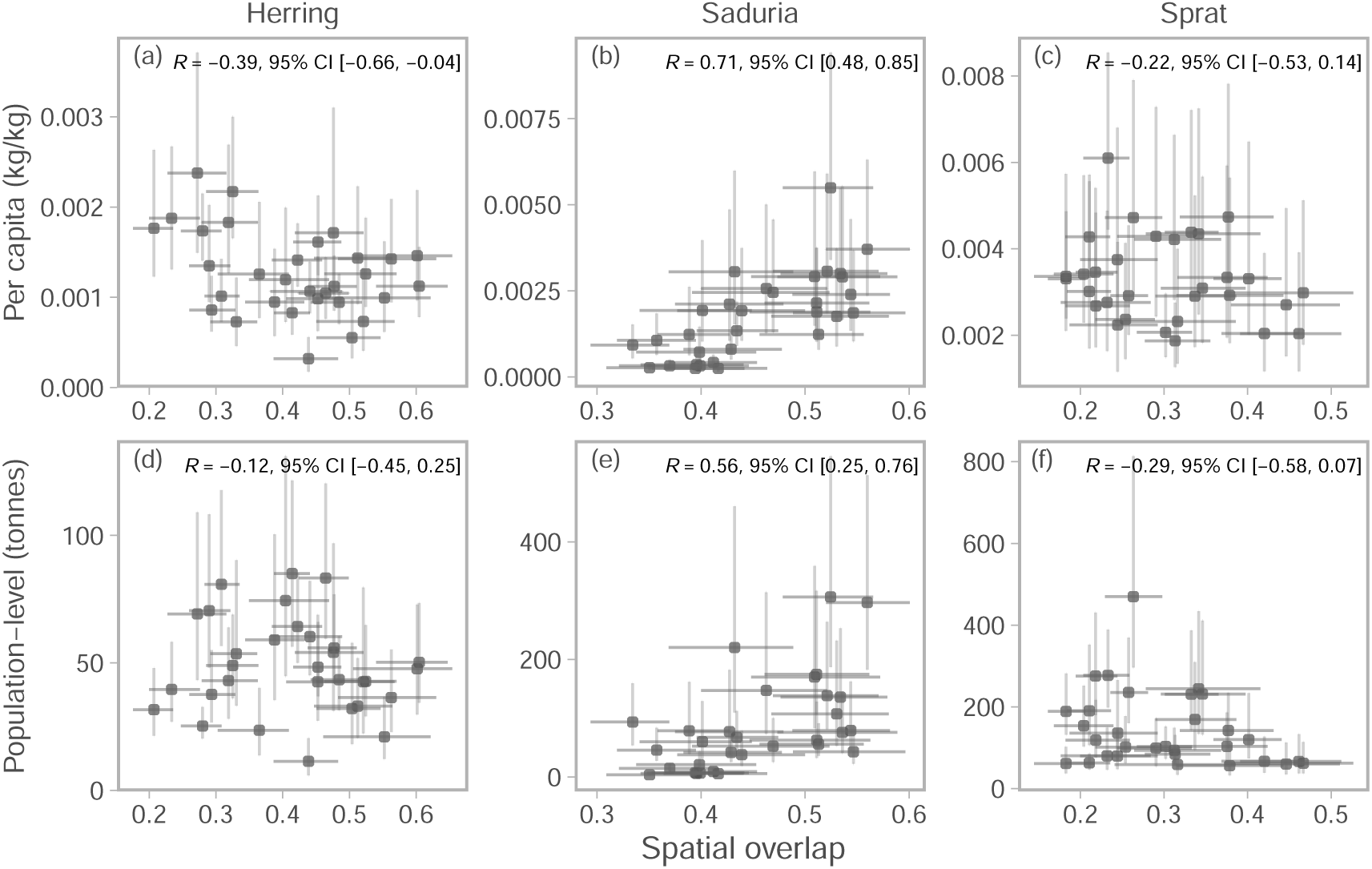
Correlation between relative per-capita predation and spatial overlap (top row, a–c), and relative population-level predation and spatial overlap (bottom row, d–f). Points depict the median, and vertical and horizontal lines depict the range between the 10^th^ and 90^th^ percentile of predation and spatial overlap, respectively, calculated from 500 simulated spatial predictions of predation and cod density. The Pearson correlation coefficient and its 95% confidence interval is printed in the top right corner of each panel for herring (a,d), *Saduria* (b, e), and sprat (c,f).

## Discussion

In this study, we quantified the effects of local prey availability on the stomach contents of predators and the relationship between predation and spatial overlap of predator and prey. Our analysis used the Baltic Sea as a case study. However, the spatiotemporal modelling approach used, which scales up local-scale diet data to population-level predation metrics while accounting for range shifts, can be applied generally, and is well suited to improve our understanding of predator-prey interactions in other systems. Our results suggest that the effects of local prey availability are modest and uncertain, and that only in the case of the benthic isopod *Saduria* availability is related to what is found in the stomachs of cod. Furthermore, only for *Saduria* do we find a correlation between predator-prey spatial overlap and per-capita and population-level predation. Our analysis therefore echoes previous studies on the challenges of linking prey dynamics to predator performance in marine generalist predators (Hilborn *et al*. 2017, Fall *et al*. 2021, Goodman *et al*. 2022), and is at contrast with the strong bottom-up effects present in trophic ecosystem models (Smith *et al*. 2011, Chagaris *et al*. 2020). Our analyses constitute important steps towards understanding the spatial scale of trophic interactions (Amarasekare 2008, Carroll *et al*. 2024), which are needed to support the implementation of ecosystem-based fisheries management.

### Effects of prey availability on feeding of generalist predators

It has been suggested that the large abundance fluctuations of many small pelagic fish species would help identifying relationships between forage fish and predators (Hilborn *et al*. 2017) because it results in large contrasts for analysis. This is also the case in our system. The Baltic Sea sprat stock increased 5-fold between 1991–1997 (ICES 2022b), albeit in the entire Baltic Sea, and not necessarily in the entire distributional range of cod. However, we did not detect an effect of sprat availability on the relative weight of sprat in cod stomachs, and the per-capita and population-level predation on sprat by cod correlated negatively (but not significantly) with cod-sprat spatial overlap. One potential reason for this could be that cod are not limited by sprat biomass but by their digestive capacity. This could limit predation even if the population-level prey biomass is low, as long as there is relatively high prey density locally (Hilborn *et al*. 2017, Fall and Fiksen 2020). Alternatively, it may be that the prey availability is not modelled at the appropriate spatial or temporal scale. Prey biomass is aggregated over ICES rectangles and is derived from a hydroacoustic survey conducted in the fourth quarter (while approximately 60% of stomach data are sampled in the fourth quarter). Hence, the spatiotemporal mismatch between data might be too large to accurately reflect predator-prey interactions, given the patchy distribution of schooling prey and the fact that stomach content data reflect consumption over a relatively short time period. Moreover, we measure overlap and pelagic prey availability in two dimensions, which may not accurately reflect encounter rates in a three dimensional environment. In line with this, Fall *et al*. (2021), also found weak effects of prey (capelin, *Mallotus villosus*) abundance on the consumption of capelin by cod. Instead they found that proximity of capelin to the seafloor was a better predictor of capelin in the cod diet, which illustrates the potential importance of the third dimension (i.e., depth). Future studies could explore whether this is also the case in the Baltic Sea, potentially using high-resolution data on schooling fish in combination with stomach content data to try and determine appropriate scales for analysis.

The spatial and temporal scale is not only relevant for the actual interaction (encounter and predation), but also when it comes to “scaling-up” functional responses or predation metrics from experimental or local scales to scales more relevant for management, e.g., to the stock- or population-level (Hunsicker *et al*. 2011). This is inherently difficult, as the relationship between consumption and prey density can change over spatial scales (Bergström and Englund 2004). Another approach, which overcomes this issue, is to use stock-level estimates when estimating functional responses (Essington and Hansson 2004). However, a limitation is that spatial heterogeneity in feeding dynamics or predator distribution and range shifts would not be accounted for. In the alternative model-based approach used here, we arrive at population-level predation metrics by summing spatially explicit predictions over the heterogeneous domain. This has the benefit that local scale heterogeneity in prey availability and stomach contents are explicitly considered, providing more accurate estimates.

### Implications for Baltic Sea cod

Our results on the spatiotemporal dynamics of predation and overlap provide novel insights that both corroborate and contrast previous hypotheses on Baltic cod. Neuenfeldt *et al*. (2020) identified that cod feeding rates on *Saduria* and sprat were substantially higher in the period 1963–1989 than in 1994–2014. Within the second time period, the physiological condition of cod declined rapidly (Mion *et al*. 2021, Eero *et al*. 2023, Lindmark *et al*. 2023). This makes it an important time period to analyze for understanding the relationship between feeding dynamics and predator-prey overlap. We observe that feeding on *Saduria* continued to decline after 2014 to near zero in the most recent years. Although the spatial overlap with *Saduria* also declined slightly, it seems that the loss of predation on *Saduria* is mainly driven by changes in per-capita predation, because trends are similar for per-capita (where the total cod biomass is factored out) and population-level predation rates. This could in turn be due to declining local abundances of *Saduria*, because even though *Saduria* does not seem to have declined substantially over the last 30 years in shallow areas (Svedäng *et al*. 2022), its area occupied may have declined, since its depth distribution is linked to oxygen dynamics on the sea floor (Karlson *et al*. 2002). This, however, is not currently possible to investigate further due to the low spatiotemporal resolution of *Saduria* biomass data. Increased competition for *Saduria* with flounder (*Platichthys* spp.) may potentially explain the declines in availability of *Saduria* to cod, as hypothesized in Haase *et al*. (2020). Support for this hypothesis is found in recent years, as high flounder density is associated with lower levels of Saduria in predator stomachs (Lindmark *et al*. 2023). However, whether competition explains the long term decline in *Saduria* in cod stomachs is not as clear, since in the 1980’s and early 1990’s (Orio *et al*. 2017), flounder was more numerous than now and cod still fed on Saduria in high numbers.

Previous studies have hypothesized that the decline in feeding rates on sprat is linked to availability and spatial overlap (Eero *et al*. 2012, Casini *et al*. 2016, Neuenfeldt *et al*. 2020), although this has never been tested explicitly. While there is a spatial mismatch in the sense that sprat biomass is highest in the northeast (Appendix S1: Figure S8c) and cod is mainly found in the southern parts, that does not necessarily mean cod are limited by low sprat abundances in the south. In line with Eero *et al*. (2012), our results suggest that predation pressure on sprat is highest in the south. We find fluctuations in the spatial overlap with sprat, with a positive trend for the per-capita predation. Also considering the lack of effect of sprat biomass on the relative weight of sprat in cod stomachs (Table 1), our study does not support the hypothesis that spatial dynamics and spatiotemporal mismatch in overlap explains trends in cod feeding on sprat. The lack of correlation between overlap and predation is also found in herring, and per-capita predation on herring by cod even increased since 2010 when the spatial overlap started to decline.

### Limitations and areas of future research

We made several simplifying assumptions that warrant future research. When calculating the predation indices, we used the total cod density in the survey catches, i.e. not resolved by length class. Since cod undergo strong ontogenetic diet shifts (Kulatska *et al*. 2019, Lindmark *et al*. 2025), it may be more appropriate to use the biomass density of cod within specific size groups that mainly feed on a specific prey. However, that may also not have a strong impact, since the correlation between local cod densities of different size groups is quite high (Jacobsen *et al*. 2023), the population size structure is highly truncated, and the main sizes currently in the population feed on a mix of pelagic and benthic prey (Lindmark *et al*. 2025). Moreover, the predation indices calculated here are only for cod of 33 cm (for simplicity, the mean size in the diet data), although they could be predicted for any size since length is a covariate in the models. This simplifying assumption effectively assumes the entire cod stock is of a specific size, which can be misleading for population-level predation metrics since the size-distribution of the cod population has been truncated over time (Eero *et al*. 2023). However, it does facilitate comparison over time for the per-capita predation metrics. Moreover, the indices are relative since the biomass density of cod is relative (due to unknown survey catchability), and since length is only a fixed effect in the model, predicting for different lengths would just change the relative value of the index identically over time.

The exact values of the predation indices are not directly comparable to mortality rates because the units are different. However, it would be straightforward to expand our work by converting our predation metrics to predation rates using gastric evacuation models (as in e.g., Gartland and Latour 2024). A qualitative comparison between our predation estimates and predation estimates from the Baltic Stochastic Multispecies Model (SMS) model (Lewy and Vinther 2004) suggests that population-level predation intensity and natural mortality of sprat and herring both increased from 1993 to until around 2010, after which cod predation declined to levels comparable to around year 2000 (ICES 2019). This implies that the stomach content data, more than model type (spatiotemporal index or multispecies assessment model) drives the result, although more work is needed to identify which processes cause discrepancies between the time series of predation and mortality. Stomach content sampling is often affected by gaps and inconsistencies in space and time (Figure 1). This represents a challenge for using stomach data in population dynamic models without some form of standardization, since there are clear spatiotemporal patterns in stomach content data. This is, however, not routinely done (ICES 2019, Neuenfeldt *et al*. 2020). The approach used in this study represents a model-based approach to estimate trophodynamic indices over the spatial domain using spatial and spatiotemporal random effects (Thorson *et al*. 2015, Cao *et al*. 2017, Karp *et al*. 2025). This approach has benefits over design-based indices by being able to include covariates and latent variables, which can improve estimates when the sampling is spatially or temporally unbalanced as the Baltic diet data are (Figure 1). This means estimates in areas with low sampling intensity for a given year depend on the ability to estimate constant and time-varying spatial random effects, which illustrates the importance of critically evaluating models (Yalcin *et al*. 2023).

We believe there are several possibilities for future work. For instance, this could include further exploration of density dependence, or at which spatiotemporal scale covariates should be included (Fall *et al*. 2021, Lindmark *et al*. 2024). Another area of research could be the covariation between different prey groups, since they are likely not independent. For instance, if cod recently fed on herring, they may not feed on benthic prey soon after, or cod may switch to sprat if the abundance of herring declines. Questions such as these could potentially be addressed using dynamic structural equation models including simultaneous or lagged effects (Thorson *et al*. 2024).

### Conclusion

Predator-prey interactions play an important role in defining ecosystem functioning and the trophic structure of marine food webs. The relationship between predator feeding dynamics and prey is a crucial aspect of this. Understanding these interactions is critical for supporting the implementation of broader food web considerations in an ecosystem-based approach to fisheries management. Essentially, this means being able to predict the ecological effects on predators that may stem from changes in prey abundance and distribution, fisheries and other anthropogenic pressures. For increases in prey to have a causal impact on predator productivity, there has to be a link between prey availability and predation. While our analysis does not control for all potential confounding factors, we do not find evidence of such positive associations between prey and predator feeding dynamics despite large fluctuations in abundance, local-scale availability, and population-level spatiotemporal overlap. While we do acknowledge that spatiotemporal dynamics of these interactions are complex and scale-dependent and may be difficult to quantify (e.g., Fall *et al*. 2021), our results could mean that the effects of specific prey are weaker than previously thought. However, it is difficult to make general statements given the mixed results in the literature (e.g., Free *et al*. 2021, Goodman *et al*. 2022), and since it may have implications for assessment and management, it should be evaluated case-by-case. In the example of Baltic cod, it could mean that management interventions aimed at increasing the availability of pelagic prey would have limited impacts on the productivity of cod.

## Acknowledgments

In memory of our dear friend and colleague Dr. Didzis Ustups, a key contributor to the manuscript, the ICES community, and to fisheries ecology in the Baltic Sea. We thank everyone involved in the collection, processing, and collation of stomach content data over time that led to the creation of the ICES stomach content database, Ivo Šics and everyone else involved in the EASME/EMFF/2018/011 SC10, Olavi Kaljuste for providing pelagic data, Hagen Radtke and Ivan Kuznetsov for assistance in acquiring predictions of *Saduria* densities, and Gemma Carroll for help with spatial overlap metrics. We also thank Robert Thorpe for a review of the paper. This work builds on the findings of the specific contract CINEA/EMFAF/2021/3.1.4-LOT1/SC10/SI2.877930*, requested by the European Climate, Infrastructure and Environment Executive Agency (CINEA), and funded through the European Maritime, Aquaculture and Fisheries Fund (EMFAF). Neither CINEA or the European Commission can guarantee the accuracy of the scientific data/information, the information does not necessarily reflect their official opinion, and they are not to be held accountable for the contents herein.

## Author contributions

Conceptualization: M.L., C.A.G., V.B., F.M.; Data curation: V.T., M.L; Methodology: M.L., C.A.G., V.B., F.M., S.C.A; Formal analysis: M.L., V.T., S.C.A; Visualization: M.L., S.C.A; Software: S.C.A; Project administration: N.S.J., M.L., C.A.G., V.B.; Resources: K.N.A., J.P., M.A., M.R., K.W., M.G., I.S., D.U.; Writing – original draft: M.L.; Writing – review & editing: All authors.

