## Supporting Information for "Weak effects of local prey density and spatial overlap on predation intensity in a temperate marine ecosystem"

### Supporting Information S1

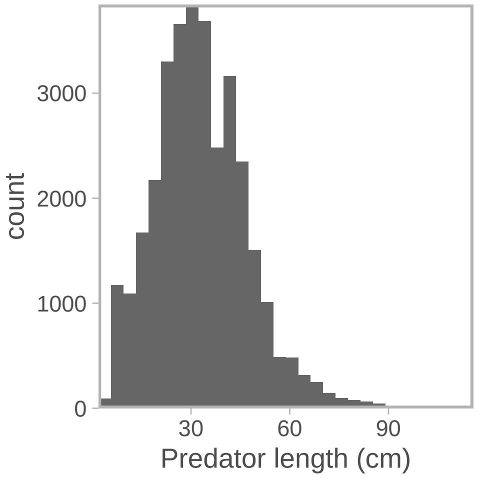


Figure S1: Distribution of cod sizes in the stomach content data.

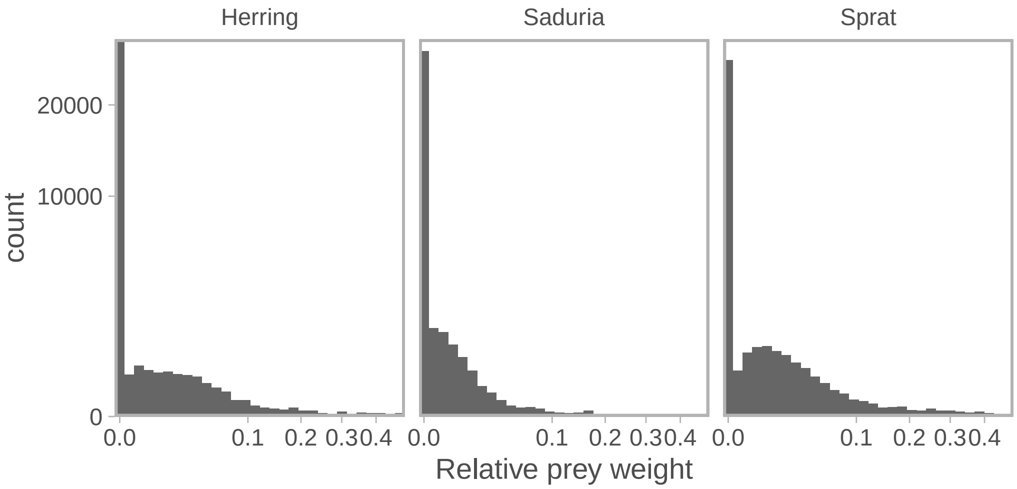


Figure S2: Distribution of relative prey weights. Note the axes are square-root transformed to better view the distribution.

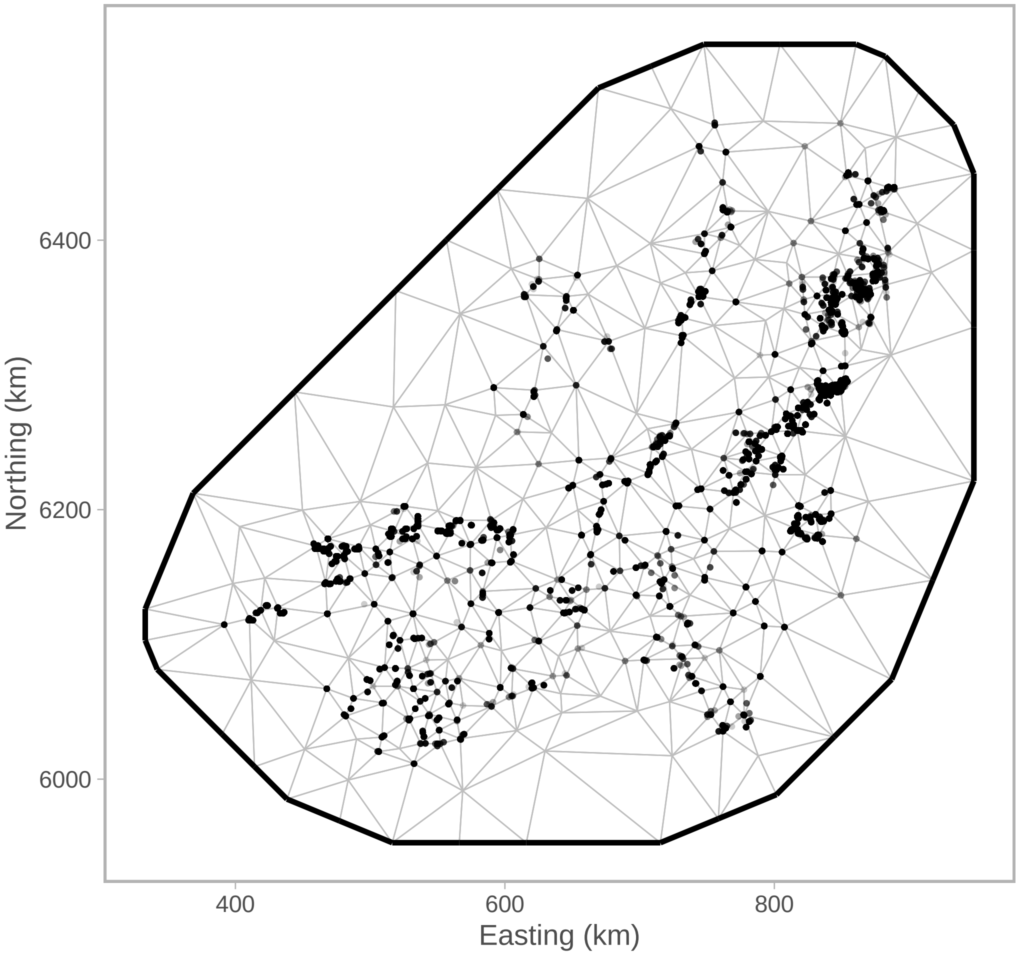


Figure S3: SPDE mesh for the stomach content models.

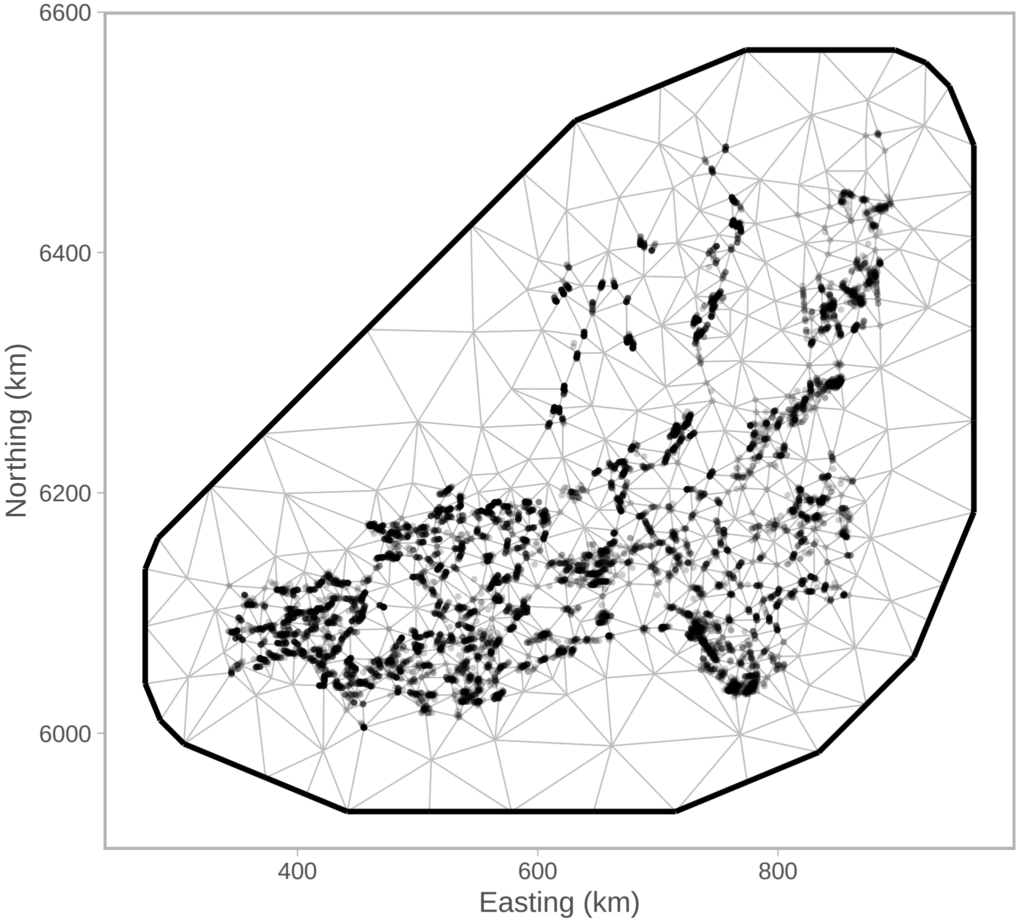


Figure S4: SPDE mesh for the biomass density model.

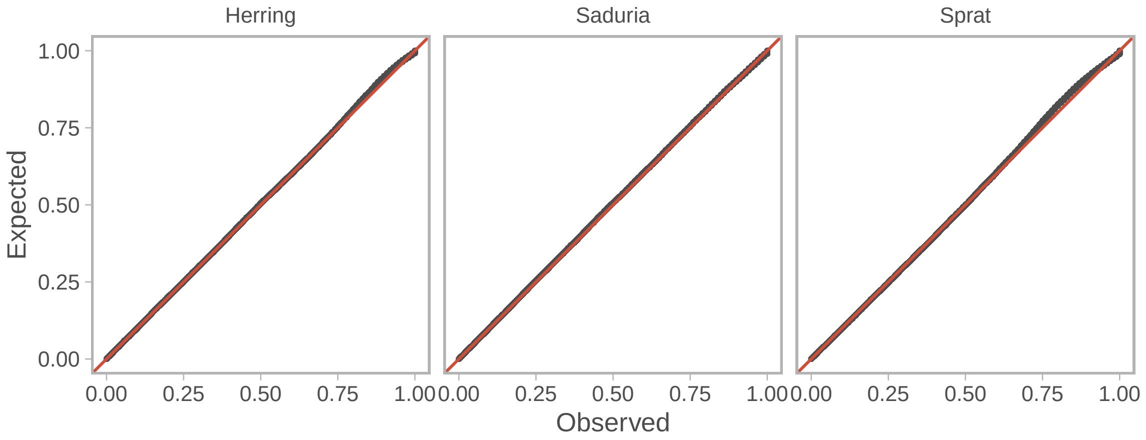


Figure S5: QQ-plots of the stomach content models based on simulated randomized quantile residuals (Dunn *et al.* 1996, Hartig 2022).

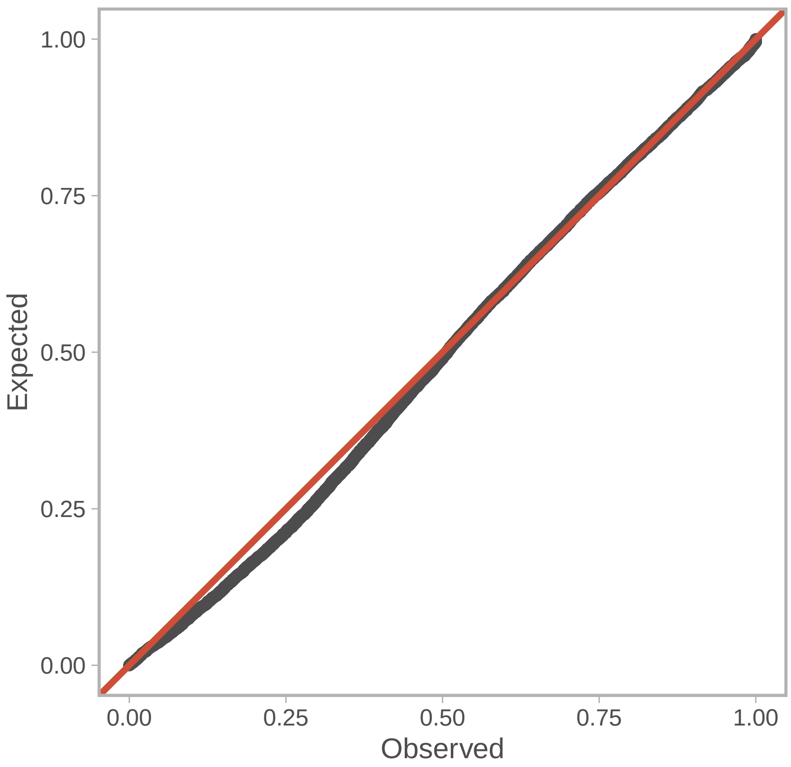


Figure S6: QQ-plots of the cod biomass density model based on simulated randomized quantile residuals (Dunn *et al.* 1996, Hartig 2022).

Table S1: Estimates for the prey-effects on the link scale (log for both linear predictors) for *Saduria*.

| **Model** | **Coefficient** | **Estimate** | **Standard error** |
|---|---|---|---|
| Binomial | Slope | 1.28 | 0.19 |
| Binomial | Breakpoint | -0.29 | 0.08 |
| Gamma | Slope | 0.25 | 0.17 |
| Gamma | Breakpoint | -0.87 | 0.36 |

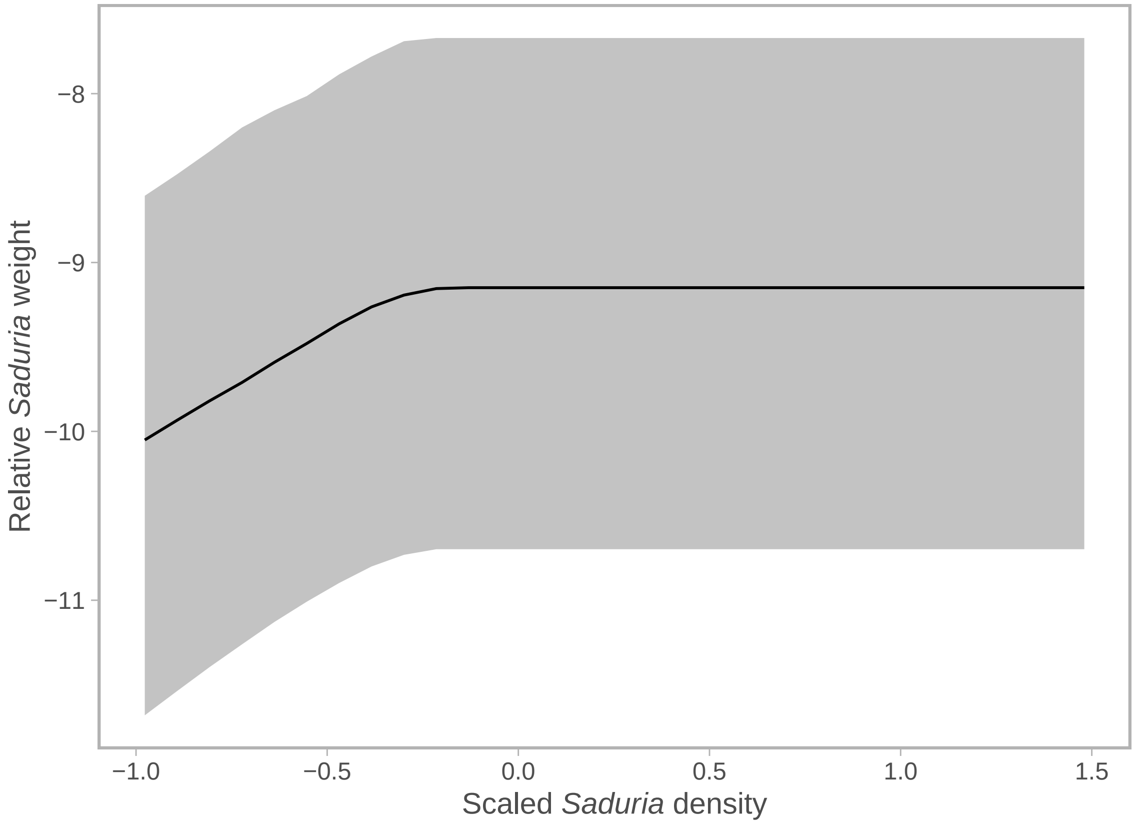


Figure S7: Conditional effects of *Saduria* biomass density for the relative prey weight of *Saduria*. The prediction (on the link scale) is for the total relative prey weight, i.e. both model components. The solid line depicts the median, and the shaded area covers the $10^{th}$–$90^{th}$ percentile of 500 draws from the joint precision matrix. The prediction is done with random effects omitted, for a cod of mean length (33 cm), and at the mean depth in 2019.

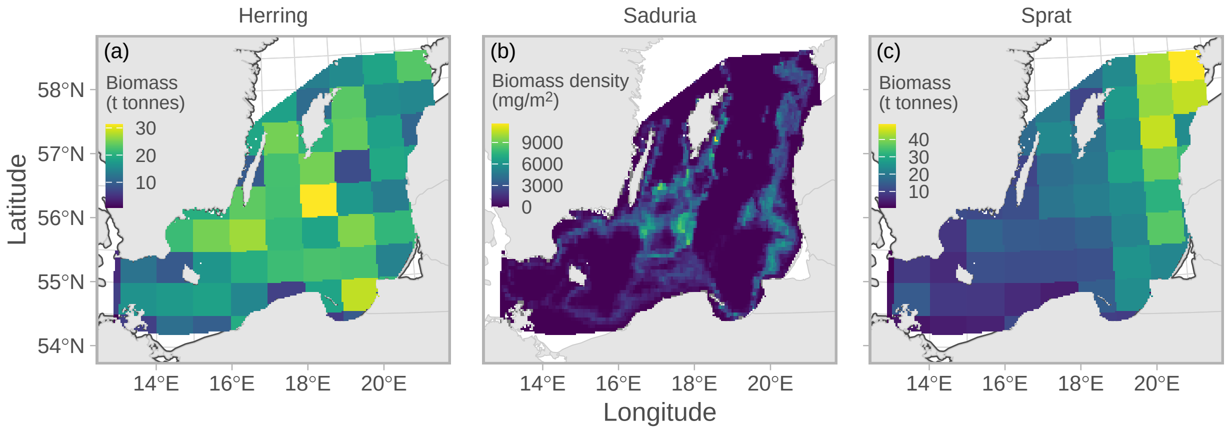


Figure S8: Prey biomass (sprat and herring) and biomass density (*Saduria*) averaged over time (1993–2023).

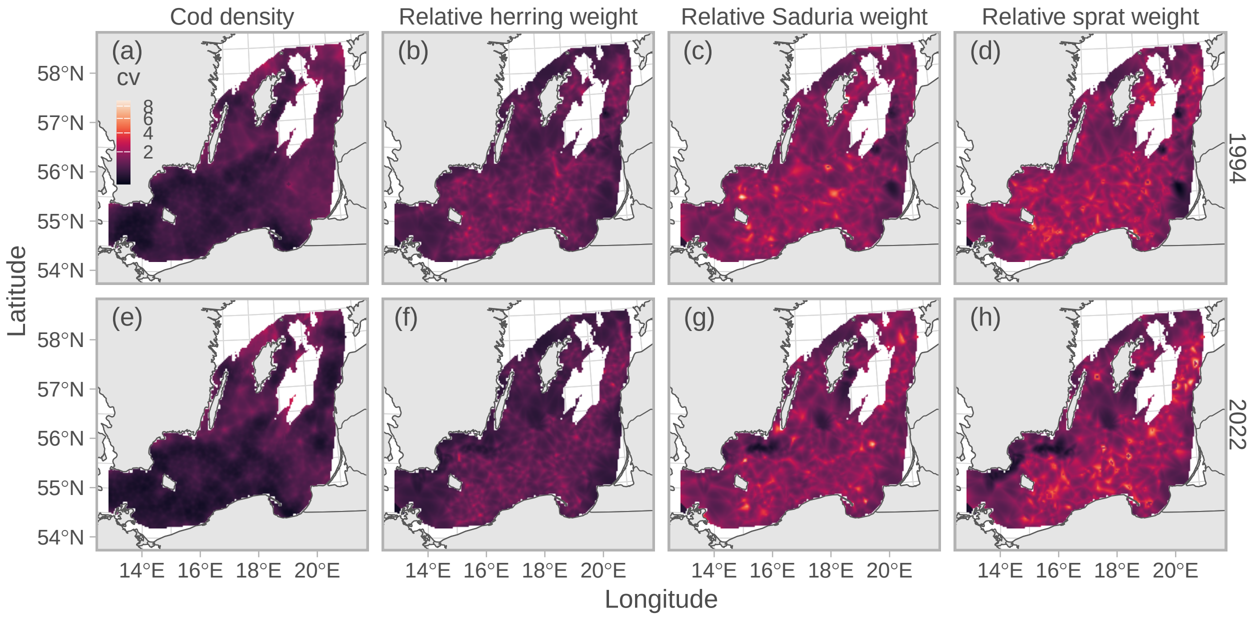


Figure S9: Coefficient of variation (CV) calculated on a grid-cell level across 500 simulations in 1994 (top row, a–d) and 2022 (bottom row, e–h), for predicted cod biomass density (a, e) and relative prey weight of herring (b, f), *Saduria* (c, g) and sprat (d, h). Note that the color scale is square-root transformed. Only grid cells with depth <130 m are included in the plot.

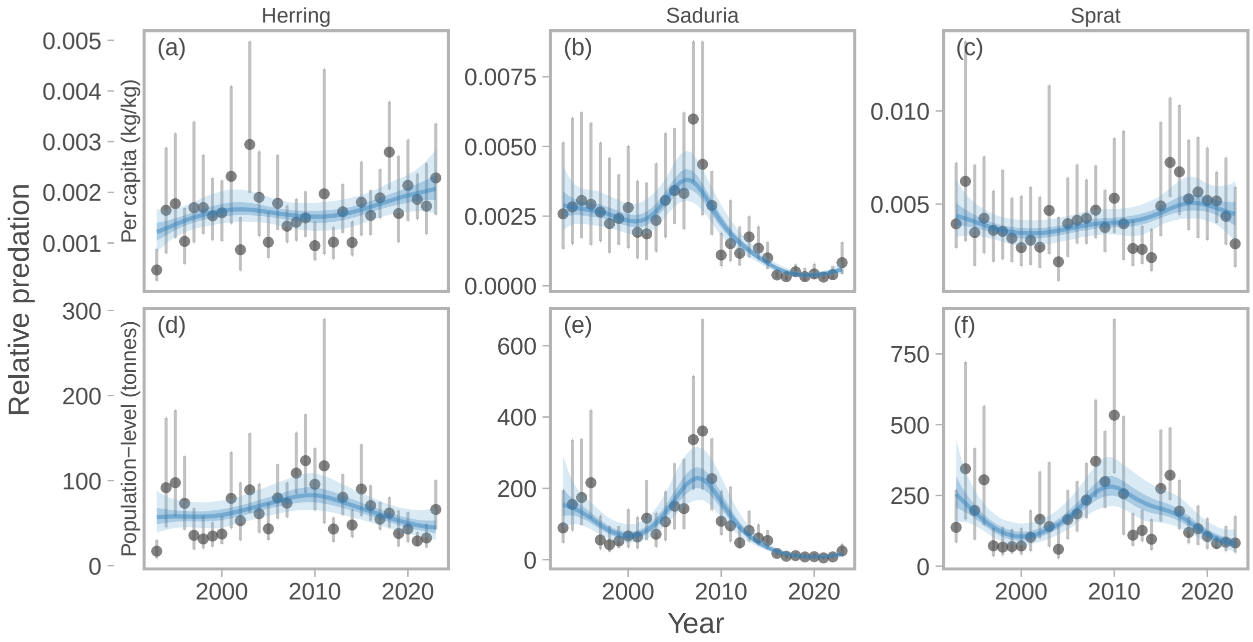


Figure S10: Relative per-capita (top row, a–c) and population-level predation (bottom row, d–f) by cod on herring (a, d), *Saduria* (b, e), and sprat (c, f) over time, *with empty stomachs omitted from the diet models*. Points depict the median predation, and vertical lines depict the range between the $10^{th}$ and $90^{th}$ percentile of predation, calculated from 500 simulated spatial predictions of both relative prey weight and cod density. Blue lines depict fits from a generalized additive model with year modelled as a penalized spline, and ribbons correspond to the 50% and 90% credible interval of the prediction.

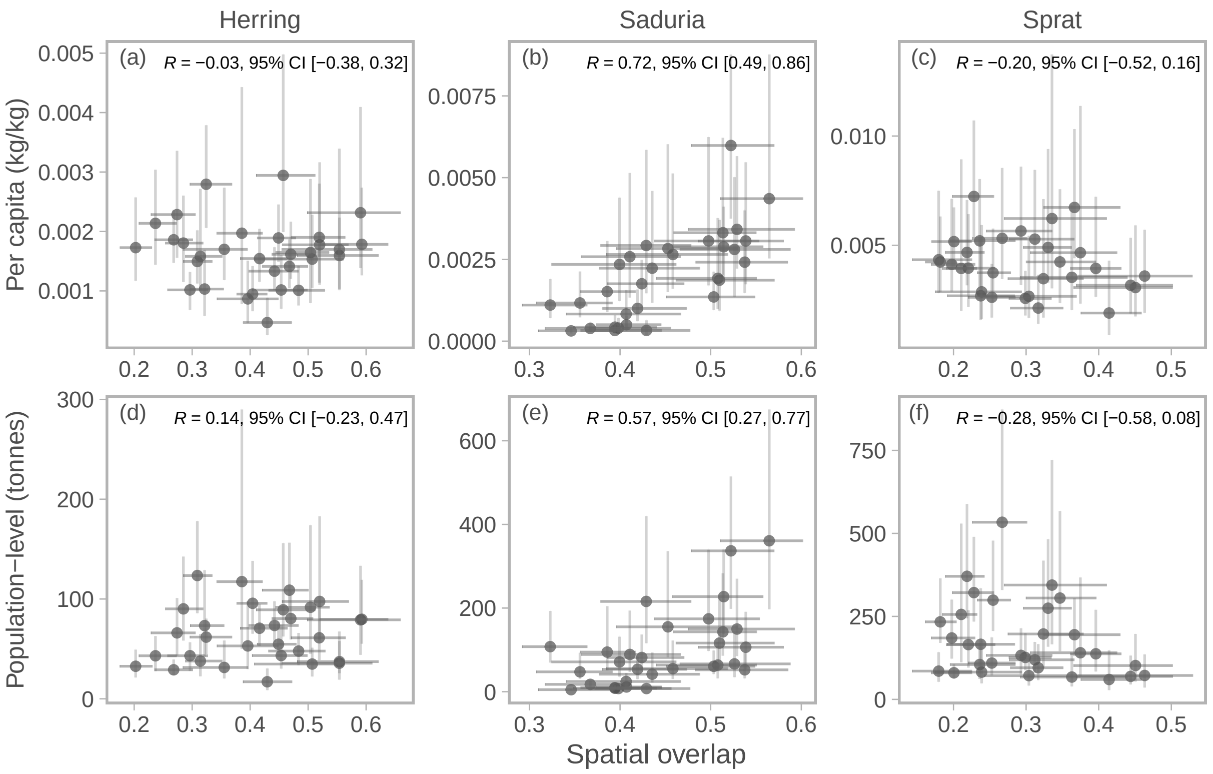


Figure S11: Correlation between relative per-capita predation and spatial overlap (top row, a–c), and relative population-level predation and spatial overlap (bottom row, d–f), *with empty stomachs omitted from the diet models*. Points depict the median, and vertical and horizontal lines depict the range between the 10$^{th}$ and 90$^{th}$ percentile of predation and spatial overlap, respectively, calculated from 500 simulated spatial predictions of predation and cod density. The Pearson correlation coefficient and its 95% confidence interval is printed in the top right corner of each panel for herring (a,d), *Saduria* (b, e), and sprat (c,f)
